# A cell-autonomous role for primary cilia in long-range commissural axon guidance

**DOI:** 10.1101/2022.08.15.503894

**Authors:** Alexandre Dumoulin, Nicole H. Wilson, Kerry L. Tucker, Esther T. Stoeckli

**Author notes:** corresponding author: E.T.S., send correspondence to: Esther T. Stoeckli.

## Abstract

Ciliopathies are characterized by the absence or dysfunction of primary cilia. Despite the fact that cognitive impairments are a common feature of ciliopathies, how cilia dysfunction affects neuronal development has not been characterized in detail. Here, we show that the primary cilium is required cell-autonomously by neurons during neural circuit formation. In particular, the primary cilium is crucial during axonal pathfinding for the switch in responsiveness of axons at a choice point, or intermediate target. Utilizing animal models and in vivo, ex vivo, as well as in vitro experiments, we provide evidence for a critical role of the primary cilium in long-range axon guidance. The primary cilium on the cell body of commissural neurons transduces long-range guidance signals sensed by growth cones navigating an intermediate target. In extension of our finding that Shh is required for the rostral turn of post-crossing commissural axons, we show here that the cilium is required for a transcriptional change of axon guidance receptors, which in turn mediate the repulsive response to floorplate-derived Shh shown by post-crossing commissural axons.

## Introduction

The primary cilium is a non-motile protrusion that localizes to the cell soma. It works as a signaling hub involved in key developmental processes, such as survival, proliferation, differentiation, polarization, and migration of cells (Goetz and Anderson, 2010). Mutations in genes that encode proteins required for primary cilia formation, maintenance, or function have a dramatic impact in human, leading to a wide spectrum of disorders, classified as ciliopathies (Reiter and Leroux, 2017). Patients with mutations in ciliary genes have a broad range of symptoms, including kidney and liver problems, limb malformations, and very often cognitive impairments (Reiter and Leroux, 2017; Valente et al., 2014). In some types of ciliopathies, like Joubert or Bardet–Biedl syndromes, the brain of patients is structurally impaired (Valente et al., 2014). For instance, axonal tracts in Joubert syndrome patients were shown to be aberrantly formed suggesting abnormal neural circuit formation and potentially axon guidance defects (Sattar and Gleeson, 2011). Therefore, we used animal models to better understand the etiology of these disorders by analyzing the role of the primary cilium during neural circuit formation.

Commissural dI1 neurons of the spinal cord have provided an informative model to study molecular mechanisms of neural circuit formation. Their axons extend ventrally from the dorsal spinal cord and cross the floorplate, the ventral midline, before turning anteriorly along the longitudinal axis. Many guidance cues for commissural axons have been identified (Chédotal, 2011; de Ramon Francàs et al., 2017; Nawabi and Castellani, 2011; Stoeckli, 2018), including Sonic hedgehog (Shh), which plays multiple roles (Zuñiga and Stoeckli, 2017). While pre-crossing axons are attracted by floorplate-derived Shh (Charron et al., 2003), axons at the midline switch their responsiveness to Shh, as post-crossing axons are repelled by Shh (Bourikas et al., 2005; Wilson and Stoeckli, 2013; Yam et al., 2012).

We have previously shown that a receptor switch for Shh is responsible for the distinct axonal behaviors (Bourikas et al., 2005; Wilson and Stoeckli, 2013; Zuñiga and Stoeckli, 2017). Boc receptors transduce an attractive response to Shh in pre-crossing axons (Okada et al., 2006) via transcription-independent signaling (Yam et al., 2009). The transient expression of Hedgehog interacting protein (Hhip) in commissural neurons, which is required for axons to turn rostrally into the longitudinal axis (Bourikas et al., 2005), is triggered by Shh itself, via a Glypican1-dependent transcriptional pathway (Wilson and Stoeckli, 2013). In turn, the presence of Hhip in the growth cone modifies the response to Shh, leading to repulsion of post-crossing axons. Thus, commissural axons encountering high levels of Shh in the floorplate might activate a transcription-dependent signaling pathway. This prompted us to test the requirement of the primary cilium for transcription-dependent Shh signaling during axon guidance, as this was shown previously for cell differentiation and patterning (Bangs and Anderson, 2017).

In recent years, the primary cilium has emerged as a critical cellular appendage for transcriptional response to Shh (Nozawa et al., 2013). Binding of Shh to Patched1 (Ptc1) promotes translocation of the Shh signaling effector Smoothened (Smo) to the cilium, where it activates Gli transcription factors (Corbit et al., 2005). In this model, ciliary function is essential for the transcriptional response to Shh. However, in axon guidance, Shh signaling was shown to be transcription-independent (Yam et al., 2009). Similarly, transcription-independent signaling was implicated in cell migration (Bijlsma et al., 2007; Brennan et al., 2012). Interestingly, the subcellular localization of Smo determines the differential responses to Shh. While Smo in the cilium activates Gli transcription factors to induce gene expression, Smo located outside the cilium favors the activation of chemotactic responses (Bijlsma et al., 2012).

Based on these findings and based on our recent results implicating the Joubert syndrome-related gene C5ORF42 (also termed CPLANE1 or JBTS17) in axon guidance in the central nervous system (CNS) (Asadollahi et al., 2018), we tested whether primary cilia were required for Shh-dependent commissural axon guidance. If attractive Shh signaling occurred via a transcription-independent pathway and the post-crossing repulsive response was triggered by cell-intrinsic mechanisms (as proposed by Yam et al., 2012), one would predict that cilia would not be required for commissural axon pathfinding. On the other hand, if floorplate-derived Shh activated the transcription of guidance genes in commissural neurons, then functional cilia might be necessary. To test these possibilities, we examined commissural axon guidance in a mouse model in which ciliary function is perturbed, and refined and extended these analyses in chicken embryos, where we could silence ciliary genes in a spatiotemporally precisely controlled manner.

Taken together, our studies indicate that primary cilia are required for commissural axon guidance in a cell-autonomous manner, allowing for a transcriptional switch in responsiveness to Shh. Our data support a model of a signaling cascade triggered at the growth cone of axons crossing the ventral midline of the CNS that involves retrograde transport of Shh to the soma. There, the signal is transduced at the primary cilium for finally inducing the transcription and expression of Hhip, the Shh receptor on post-crossing axons. These data provide evidence for a long-range axon guidance mechanism involving the primary cilium as a critical signaling component allowing axons to correctly navigate in the developing CNS.

## Results

### Commissural neurons carry a primary cilium *in vivo* before, during, and after their axons cross the midline of the central nervous system

We chose the dorsally localized commissural dI1 interneurons in the developing spinal cord as an axon guidance model. These commissural neurons provide a very accessible neuronal population to investigate molecular mechanisms of axon guidance, as they have a very stereotypical trajectory towards and across the ventral midline of the spinal cord. Their axons approach the ventral midline at the floorplate level, cross it, exit it, and turn rostrally towards the brain (Stoeckli, 2018).

We first asked whether dI1 neurons carried a primary cilium during the time window of axon pathfinding. Primary cilia were found on the dI1 subpopulation of commissural neurons in both chicken and mouse embryos (Fig. 1). We used *in ovo* electroporation of a plasmid for dI1 neuron-specific labeling (Wilson and Stoeckli, 2011) together with immunostaining of dI1 neuron nuclei (Lhx2 transcription factor) and primary cilia (Arl13b) to reveal the presence of primary cilia at different stages and to visualize the location of the dI1 axon tips at these time points (Fig. 1A-L). Arl13b-positive cilia were found on dI1 commissural neuron cell bodies in the chicken spinal cord during initial axon growth (Hamburger and Hamilton stage (HH)20), pre-crossing axonal elongation (HH22), midline crossing and exiting (HH24), and during extension of post-crossing axons along the contralateral floorplate border (HH26, white arrows point to cilia and white arrowheads to dI1 growth cones, respectively, Fig. 1A-L).

**Figure 1.**
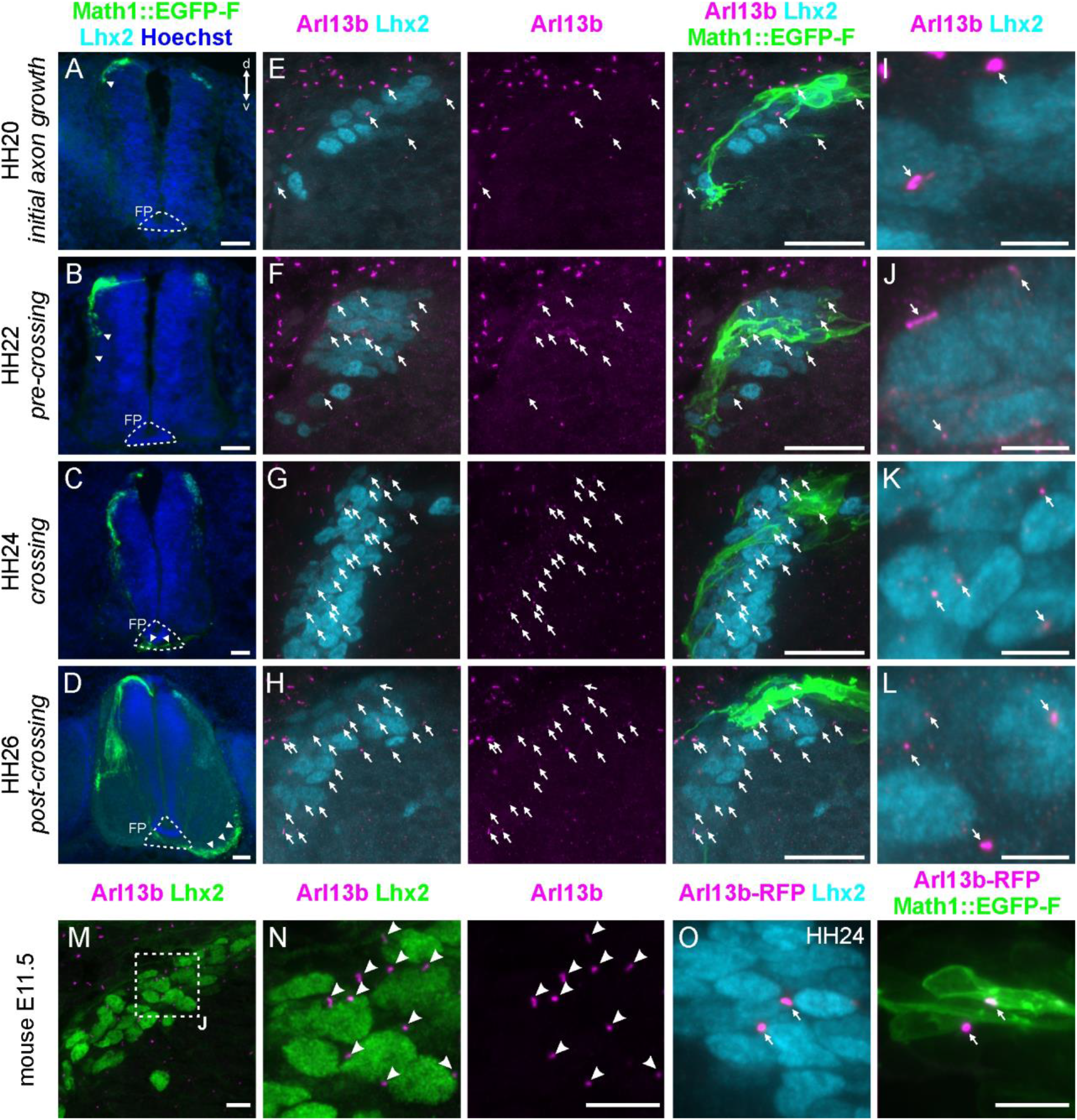
The dI1 commissural neurons carry a primary cilium at different time points of their development in vivo. (A-D) Transverse sections of chicken embryos in which the Math1::EGFP-F plasmid (green) was electroporated unilaterally to label dI1 neurons at HH17-18. Embryos were sacrificed at different time points. At HH20, dI1 axons were starting to extend an axon (A). At HH22, they were growing ventrally (B), at HH24, they were crossing the floorplate (C), and at HH26, post-crossing axons were localized in the contralateral ventral funiculus (D). Arrowheads show where dI1 axonal growth cones localized at the different time points. Lhx2 was used as a marker for dI1 nuclei (cyan) and section were counterstained with Hoechst to stain all nuclei (blue). (E-H) High magnification pictures of the dI1 neuron area of the same spinal cords depicted in (A-D) showing Lhx2-positive dI1 nuclei (cyan) co-stained with the primary cilium marker Arl13b (magenta) and GFP (dI1 neuron reporter, green). These neurons carried a primary cilium (arrows) on their soma throughout stages HH20 to HH26. (I-L) Cropped pictures of an area shown in (E-H). (M-N) Similar observations were made in E11.5 mouse embryos at the time, when dI1 axons were crossing/exiting the midline area with Lhx2-positive dI1 neurons (green) carrying Arl13b-positive primary cilia (red, arrowheads). (O) Ciliation of dI1 neurons was confirmed by co-electroporating Arl13 fused to RFP (Arl13b-RFP) and Math1::EGFP-F in vivo at HH17-18 and co-staining of RFP (magenta), GFP (green) and Lhx2 (cyan) on HH24 spinal cord sections (white arrows). FP, floorplate; d, dorsal; v, ventral. Scale bars: 50 μm (A-D), 25 μm (E-H), 5 μm (I-L) and 10 μm (M-O).

These data were in line with previous reports in the chicken spinal cord with newly differentiated interneurons *in vivo* and cultured commissural neurons, respectively (Toro-Tapia and Das, 2020; Yusifov et al., 2021). Moreover, the ciliation of dI1 neurons in the chicken was confirmed by immunostaining for adenylate cyclase III (ACIII), a known neuronal primary cilia marker (Fig. S1) (Caspary et al., 2016; Ou et al., 2009). Similarly, Arl13b-positive cilia were found on embryonic (E11.5) mouse dI1 neurons at a stage when their axons were crossing and exiting the ventral midline of the CNS (white arrowheads, Fig. 1M,N). In addition, we also overexpressed Arl13b-RFP *in vivo* to visualize primary cilia in the chicken spinal cord together with dI1-specific GFP expression and could confirm that these neurons carried a primary cilium at the time when their axons were crossing the floorplate by staining of RFP (white arrows, Fig. 1O).

Taken together, our results showed that dI1 neurons carry a primary cilium before their axons contact the floorplate, during floorplate crossing, and after their axons turn rostral into the longitudinal axis. Therefore, the presence of primary cilia on dI1 neurons is compatible with a role during axon guidance at a choice point.

### Commissural axon guidance is perturbed in the *cobblestone* mutant

To study the role of cilia and Shh signaling in commissural axon guidance, we examined a mouse mutant in which ciliary function is perturbed. *Cobblestone (cbs)* mice are hypomorphic for the intraflagellar transport protein-88 (Ift88), as they express *Ift88* mRNA and protein at only 25% of the levels of wildtype (WT) embryos (Willaredt et al., 2008). Ift88 is a component of the IFTB anterograde transport complex of the cilium, which is required for formation and maintenance of cilia and transcription-dependent Shh signaling (Bangs and Anderson, 2017; Huangfu and Anderson, 2005; Liu et al., 2005). Although *cbs* embryos still possess cilia, albeit in reduced numbers, they phenocopy perturbations in Shh pathway components, suggesting that the existing cilia do not adequately mediate Shh signaling (Gazea et al., 2016; Willaredt et al., 2008). We found that *cbs* homozygous embryos had a reduced number of cilia on dI1 neurons compared to WT littermates using co-staining of the cilia marker Arl13b and the dI1 neuron marker Lhx2 (Fig. 2A-D). Moreover, we found that *Shh* mRNA and protein were reduced or absent in the *cbs* floorplate (Fig. 2E and Fig. S2). Hence, we used *cbs* mice to assess the effects of reduced Shh levels and/or compromised transcription-dependent signaling on commissural axon guidance. In wildtype mice, as previously described in rat (Yam et al., 2012) and chick (Bourikas et al., 2005), we found that *Shh* mRNA was expressed in a posterior^high^ to anterior^low^ gradient along the floorplate (Fig. S2A-D).

**Figure 2.**
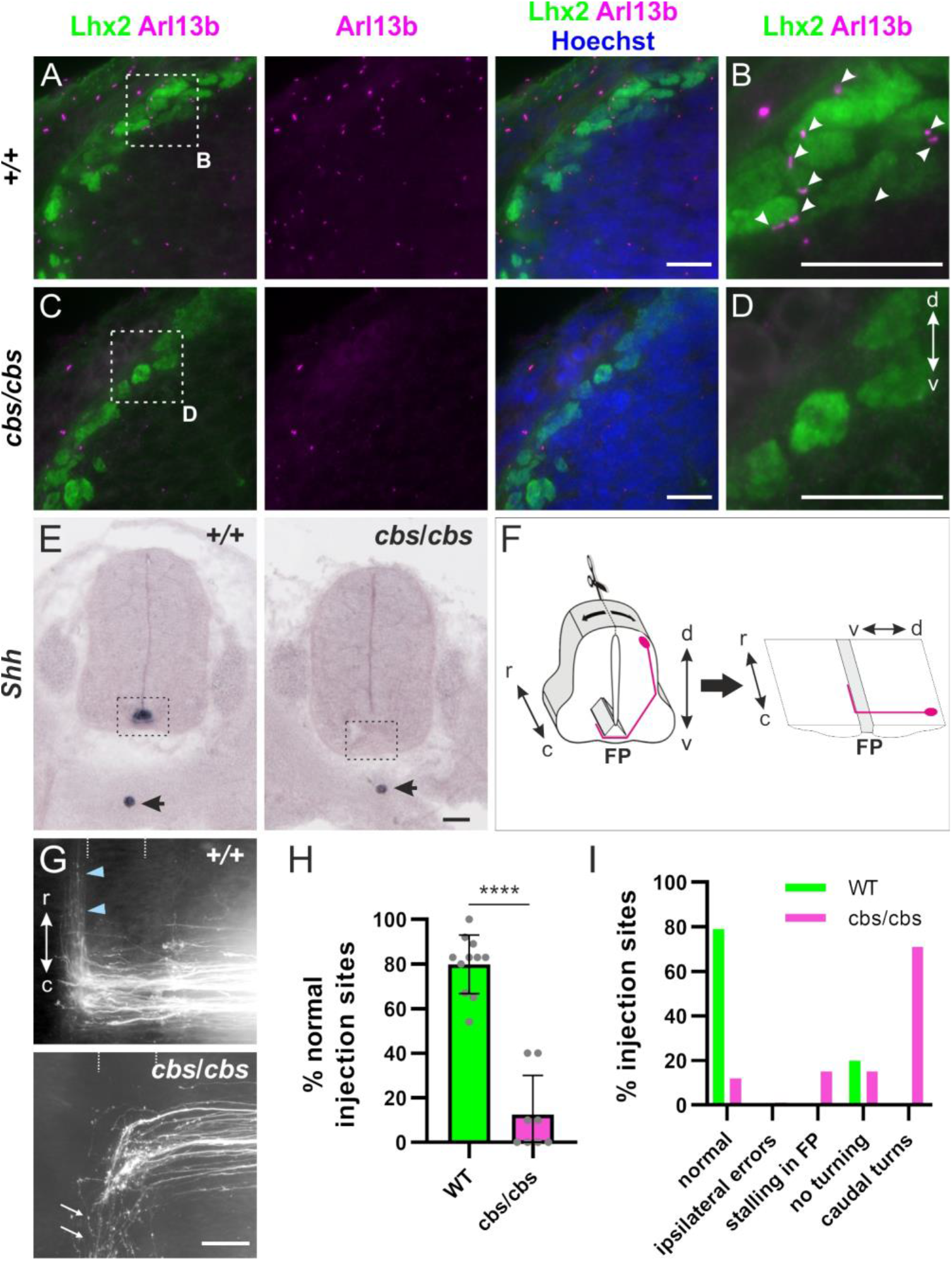
Cobblestone (cbs) mice display defects in post-crossing commissural axon guidance. (A-D) Spinal cord transverse sections of E11.5 mouse embryos were stained for the primary cilia marker Arl13b (magenta), co-stained for the dI1 neuron marker Lhx2 (green), and counterstained with Hoechst (blue). While many primary cilia were present in the area of dI1 somas of wild-type embryos (+/+, white arrowheads, B), very few to none were detected in this area in homozygous cobblestone (cbs/cbs) littermates (D). (E) In situ hybridization on E12.5 spinal cord transverse sections revealed that Shh was absent from the floorplate (dashed rectangle) in cbs/cbs embryos, whereas it was still expressed in the notochord (black arrow). (F) Schematic depicting the open-book preparation of embryonic spinal cords allowing for the visualization of dI1 axon (magenta) projections at the ventral midline, the floorplate. (G) Examples of DiI tracing of dI1 axons in an open-book preparation of a E12.5 spinal cord taken from a wild-type embryo, showing a normal rostral turning phenotype (blue arrowheads) and aberrant phenotypes seen in cbs/cbs embryos, mostly caudal turns (white arrows)). (H) Quantification of axon guidance defects in cbs/cbs embryos compared to wild-type littermates (unpaired T-test). N(embryos)= 11 (WT) and 8 (cbs/cbs); n(injection sites) = 118 (WT) and 52 (cbs/cbs). Error bars represent standard deviation. p<0.0001 (****). d, dorsal; v, ventral; r, rostral; c, caudal; FP, floorplate. Scale bars: 20 μm (A-D) and 50 μm (E and G). Source data and statistics are available in Source Data spreadsheet.

We assessed dI1 axon pathfinding by tracing axons with the lipophilic dye DiI in open-book preparations of spinal cords taken from E12.5 embryos (Fig. 2F-I). In WT or heterozygous littermates, the majority of DiI-traced axonal trajectories displayed the normal phenotype: at 80 ± 13% (mean ± standard deviation) of the DiI injection sites, axons crossed the floorplate and turned rostrally along the contralateral floorplate border (blue arrowheads, Fig 2G,H). However, in *cbs* mice, axons at only 13 ± 18% of the DiI injection sites (mean ± standard deviation) showed normal behavior. Upon reaching the exit site of the floorplate, most of the axons turned caudally instead of rostrally in cbs mice (Fig. 2G-I). Thus, perturbation of Ift88 levels caused aberrant commissural axon guidance.

### *Cbs* mice exhibit defects in ventral spinal cord patterning

Because transcription-dependent Shh signaling is required for spinal cord patterning, the observed axon guidance defects could also be caused indirectly through changes in floorplate induction and aberrant cell differentiation. Thus, we assessed whether spinal cord patterning was affected in *cbs* mice (Fig. S3). Indeed, this is what we found. At E10.5, some Islet1-positive cells erroneously invaded the ventral midline (white arrowhead, Fig.S3). The characteristic distribution of Islet1-positive motoneurons was not seen in *cbs* mice. However, by E12.5, Islet1 staining resembled WT. Nkx2.2, a Shh target that is only induced by high levels of Shh, was strongly reduced in *cbs* embryos. The few Nkx2.2-positive cells were disorganized compared to WT embryos (yellow arrows, Fig. S3). Both Shh and the floorplate marker HNF3β were absent from the *cbs* spinal cord (white arrows, Fig. S3). In contrast, dorsal markers, like Pax3, were unaffected in *cbs* mice at E10.5 and E12.5. Importantly, the differentiation of dI1 neurons still occurred normally in *cbs* mice, as Lhx2-positive differentiated interneurons could be detected (Fig. 2A-D). Moreover, these commissural dI1 neurons extended their axons normally towards the ventral midline as visualized by Axonin-1 staining (Fig. S3).

Taken together, the patterning analysis revealed that the dorsal *cbs* spinal cord was correctly specified, while in the ventral spinal cord, the floorplate and its neighboring cell types were miss-pecified.

As the floorplate is the intermediate target for commissural axons, its improper differentiation /reduction in *cbs* mice might contribute to the observed axon guidance errors. Indeed, Gli2^-/-^ mice, which lack a floorplate, display similar guidance errors to those reported here (Matise et al., 1999; Yam et al., 2012). To determine whether the axon guidance anomalies in *cbs* mice arose only secondarily to the lack of a defined floorplate, or whether Ift88 or ciliary function might also be directly required in commissural neurons for correct guidance, we next turned to *in ovo* RNAi experiments, in which we could avoid the early morphological effects of diminished Ift88 function.

### Temporally controlled loss of the IFTB proteins Ift88 and Ift52 in the chicken spinal cord leads to commissural axon guidance defects without affecting spinal cord patterning

Using *in ovo RNAi* allowed us to precisely control Ift88 knockdown in such a way that floorplate development was not affected. *Ift88* was expressed throughout the chick neural tube during commissural pathfinding (HH18-26) (Fig. S4A). Unilateral electroporation of long double-stranded RNA (dsRNA) derived from *Ift88* was performed at HH17-18 to knockdown Ift88 in one half of the neural tube only after early spinal cord patterning was completed, but just prior to commissural axon outgrowth. The efficiency and specificity of the knockdown using dsRNA was verified using a reporter assay *in vitro* (Fig. S5). After this temporally controlled silencing of Ift88, floorplate morphology and HNF3β expression were normal (Fig. S4B) and neural tube patterning was not affected, as this is completed before our experimental knockdown of Ift88 (Fig. S4C).

Silencing *Ift88* in this temporally controlled manner still caused axons to stall in the floorplate or at the contralateral floorplate border (Fig. 3A-B). Most of the axons failed to turn longitudinally, suggesting that Ift88 is directly required for correct guidance of post-crossing axons (asterisks, Fig. 3B-D). At only 36 ± 28% of the DiI injection sites (mean ± standard deviation), dI1 axons turned rostrally at floorplate exit (Fig. 3C). Importantly, embryos in which the neural tube was electroporated unilaterally with only a plasmid encoding GFP (65 ±28% of rostral turning) or the plasmid together with dsRNA against RFP (64 ± 15% of rostral turning) did not show any significant defects in axon guidance compared to untreated embryos (82 ± 10% of rostral turning, mean ± standard deviation). The majority of the axons turned rostrally at the contralateral floorplate border (blue arrowheads, Fig. 3A and Fig. 3C,D). In support of these findings, we observed similar phenotypes when we knocked down Ift52, another component of the IFTB complex that directly binds Ift88 (Lucker et al., 2010). We found that axons behaved normally at only 42 ± 15% of the DiI injection sites (mean ± standard deviation), with most of the axons showing post-crossing errors (Fig. 3C,D). Thus, Ift88 deficiency directly contributes to axon guidance errors in the chicken spinal cord.

**Figure 3.**
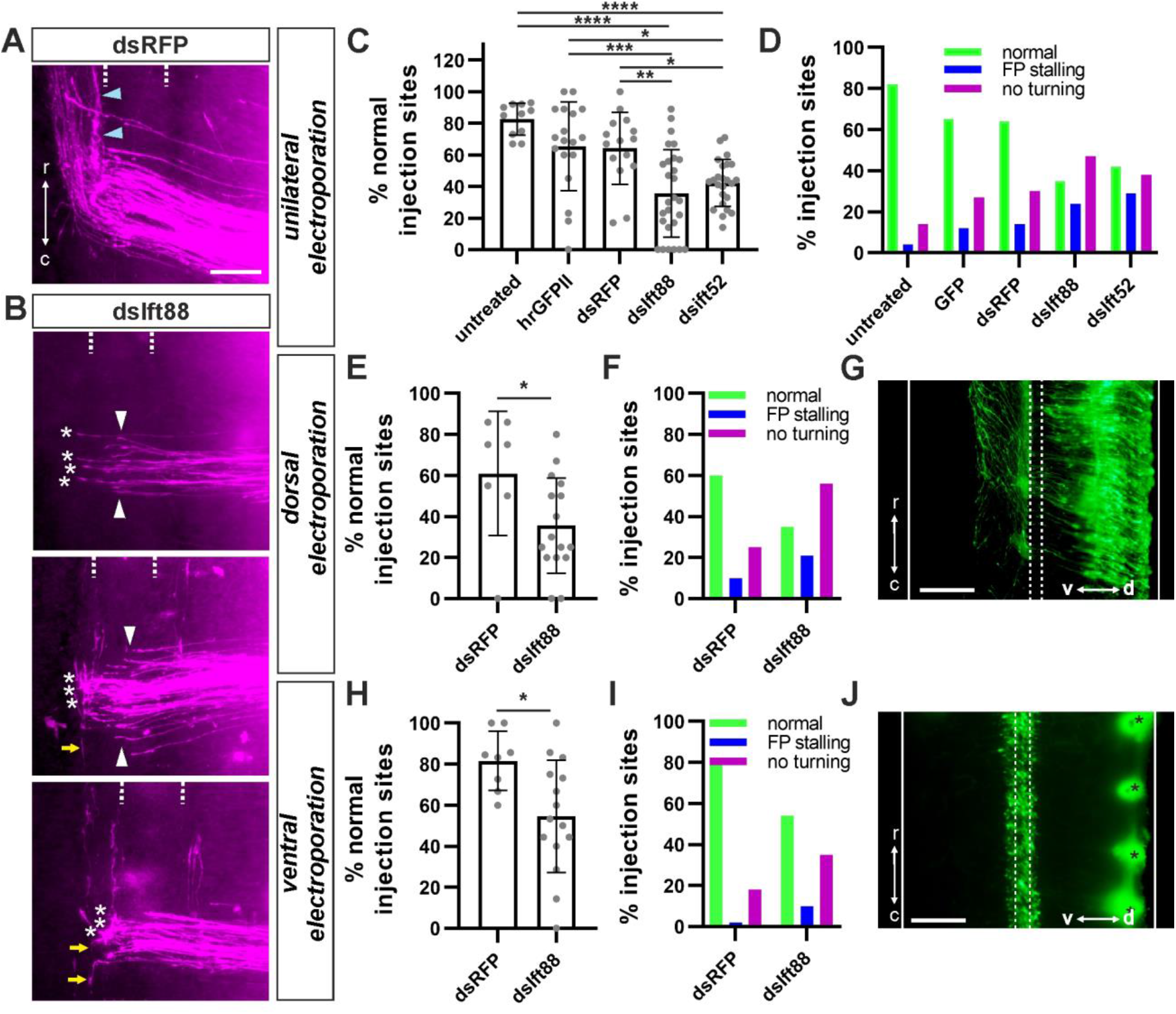
The IFTB proteins Ift88 and Ift52 are required for commissural axon guidance. (A) Representative example of DiI-traced dI1 axons in an open-book preparation of a dsRFP-injected control embryo, showing normal rostral turning (blue arrowheads). (B) Ift88 knockdown with dsIft88 induced aberrant phenotypes at the CNS midline with most of the axons being unable to turn rostrally and instead stalling (white asterisks) or turning caudally (yellow arrows) at the floorplate exit site. Moreover, some axons stalled in the floorplate area (white arrowheads). (C,D) Quantification of axon guidance phenotypes at the spinal cord ventral midline after unilateral electroporation of the spinal cord. One-way ANOVA with Tukey’s multiple comparisons test. N(embryos)= 11 (untreated), 18 (hrGFPII), 15 (dsRFP), 27 (dsIFT88), 24 (dsIft52); n(injection sites)= 135 (untreated), 178 (hrGFPII), 141 (dsRFP), 251 (dsIFT88), 323 (dsIft52). (E,F) Dorsal electroporation of dsIft88, targeting commissural neurons, induced similar aberrant phenotypes as seen after unilateral electroporation. Unpaired T-test. N(embryos)= 7 (dsRFP) and 16 (dsIft88); n(injection sites)= 49 (dsRFP) and 90 (dsIft88). (G) Dorsal targeting was confirmed by expression of co-electroporated hrGFPII (green) in open-book preparations. (H,I) Ventral electroporation of dsIft88 targeting the ventral midline also affected dI1 commissural axon guidance. Unpaired T-test. N(embryos)= 8 (dsRFP) and 15 (dsIft88); n(injection sites)= 63 (dsRFP) and 130 (dsIft88). (J) Ventral targeting was confirmed by expression of co-electroporated hrGFPII (green) in open-book preparations. Asterisks indicate DiI injection sites in this preparation (bleed-through fluorescence). Dashed lines represent the floorplate boundaries. p<0.0001 (****), p<0.001 (***), p<0.01 (**), p<0.05 (*) and p>0.05 (ns). r, rostral; c, caudal; d, dorsal; v, ventral; FP, floorplate. Source data and statistics are available in Source Data spreadsheet.

### Ift88 is required in commissural neurons for correct axon guidance at the CNS midline

To dissect the cell type-specific requirement for Ift88, we used targeted electroporation to knockdown Ift88 either in the dorsal spinal cord (targeting primarily commissural neurons; Fig. 3E,F and G) or in the ventral spinal cord (targeting the floorplate; Fig. 3H,I and J). The ventral downregulation of Ift88 mildly affected commissural axon guidance (55 ± 27% of DiI sites with rostral turning) compared to the electroporation of dsRNA targeting RFP as a control (82 ± 15% of DiI sites with rostral turning, mean ± standard deviation, Fig. 3H,I). In contrast, dorsal targeting of dsIft88 resulted in similar guidance defects to those observed after unilateral electroporation with most of the axons failing to turn rostrally (36 ± 22% of DiI sites with rostral turning) in the longitudinal axis compared to dsRFP control (61 ± 30% of DiI sites with rostral turning, mean ± standard deviation, Fig. 3E,F).

Together, the spatiotemporal knock-down of the ciliary gene *Ift88* indicated that Ift88 was required cell-autonomously in commissural neurons for correct axon guidance and that this activity was distinct from its earlier role in floorplate morphogenesis and cell differentiation (Goetz and Anderson, 2010; Tasouri and Tucker, 2011). It also suggested that a functional primary cilium is required in commissural neurons for correct axon guidance at an intermediate target.

### Ift88 is required for the transcriptional switch of Shh receptors in commissural neurons

We previously identified a Shh-mediated transcriptional switch of axon guidance receptors as responsible for the change from attraction to repulsion between pre- and post-crossing commissural axons (Wilson and Stoeckli, 2013). This transcriptional switch, leading to the transient expression of the Shh receptor Hhip, is triggered by Shh itself via Glypican-1 (Wilson and Stoeckli, 2013). Interestingly, the phenotypes reported above after silencing Ift88 were similar to those seen after the perturbation of Shh, Glypican-1, or Hhip function (Bourikas et al., 2005; Wilson and Stoeckli, 2013). Based on these results, we hypothesized that the axon misprojections observed after *Ift88* silencing could be caused by the absence of the transient expression of *Hhip* in dI1 neurons. To investigate this idea, we performed *in situ* hybridization for *Hhip* in HH25 spinal cord transverse sections after unilaterally silencing *Ift88 in vivo.* Indeed, when Ift88 was unilaterally knocked down at HH18, *Hhip* signal intensity in the dI1 neurons located in dorsal spinal cord (white dashed circles, Fig. 4) at HH25 was lower on the electroporated side (Fig. 4B). As expected, the unilateral electroporation of a plasmid encoding a GFP protein alone (control) did not lead to a reduction of *Hhip* expression in dI1 neurons (white dashed circles, Fig. 4A). Calculation of a normalized *Hhip* intensity ratio in dI1 neurons revealed that silencing *Ift88* significantly reduced the expression of *Hhip* by about 26% compared to the electroporated side with a ratio^electro:control^ of 0.74±0.19 compared to 1±0.14 for the control sample (mean ± standard deviation, Fig. 4C; N=5 embryos each, p<0.05). The reduction in Hhip expression can explain the failure of post-crossing commissural axons to turn rostrally (Bourikas et al., 2005; Wilson and Stoeckli, 2013).

**Figure 4.**
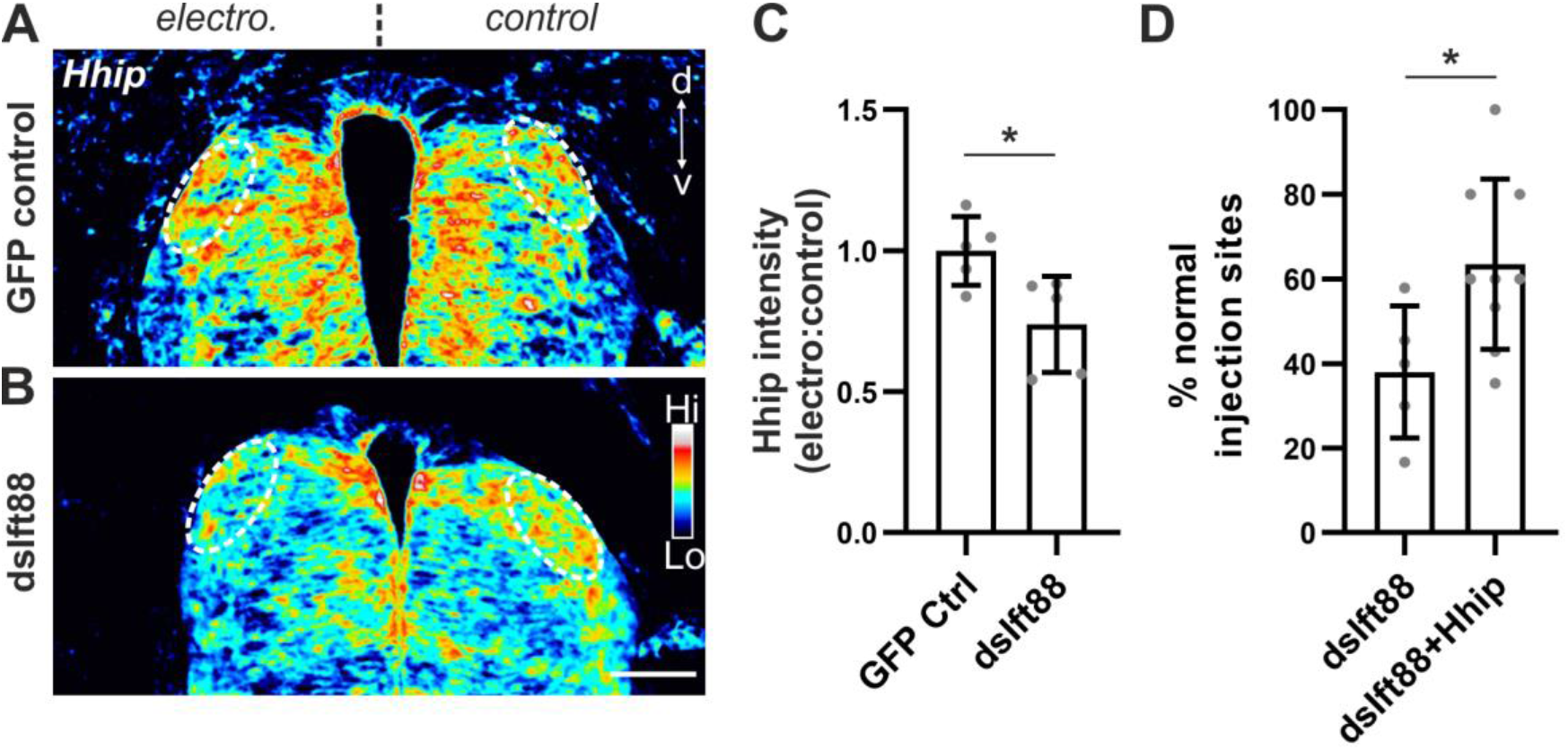
Ift88 is required for the transcription of Hhip, the Shh receptor for post-crossing commissural axons. (A-B) Knockdown of Ift88 at HH18 reduced Hhip mRNA expression in dI1 neurons at HH25. (A) Heatmap images of in situ hybridization for Hhip in the spinal cord revealed no apparent change in Hhip mRNA in dI1 neurons area (dashed ovals) when GFP was electroporated unilaterally. (B) However, co-electroporation of dsIft88 reduced Hhip mRNA expression in dI1 neurons on the electroporated side. (C) Quantification of the average Hhip mRNA intensity ratio between electroporated and control side revealed a reduction of about 25% after Ift88 knockdown compared to GFP control (unpaired T-test). N(embryos)=5 for each condition. (D) Axon guidance errors seen after downregulation of Ift88 were rescued by expression of Hhip. The number of DiI injection sites with normal axonal trajectories were significantly increased by Hhip expression compared to Ift88 loss of function and was rescued to a level similar to GFP controls (see Fig. 3C, unpaired T-test). N(embryos)= 5(dsIft88) and 9(dsIft88+Hhip); n(injection sites)= 62 (dsIft88) and 79 (dsIft88+Hhip). Error bars represent standard deviation. p<0.05 (*). electro, electroporated; d, dorsal; v, ventral; Hi, high; Lo, low. Source data and statistics are available in Source Data spreadsheet.

To provide evidence further supporting our model that Ift88 function is required for Shh-mediated induction of *Hhip* expression, and thus, repulsion of post-crossing axons, we carried out rescue experiments. To this end, we expressed mouse Hhip in dI1 neurons lacking Ift88. Consistent with our hypothesis, restoring Hhip expression resulted in normal axon guidance. We found normal axonal navigation at 63.4±6.7% of the DiI injection sites (mean ± standard deviation; Fig. 4D). This was not significantly different from control-treated embryos (see Fig. 3C). In comparison, after knocking down Ift88 we found normal axon guidance at only 38.0±6.9% of the injection sites (mean ± standard deviation, p<0.05).

Taken together, the reduction of *Hhip* expression in dI1 neurons after Ift88 knockdown and the rescue of the Ift88-dependent axon guidance errors by rescuing Hhip expression support the idea that Ift88 is part of the Shh-Glypican1-Hhip signaling cascade by acting upstream of Hhip. Our results also suggest that a functional primary cilium is required for proper induction of Hhip transcription in these neurons.

### Live imaging of dI1 commissural axons *ex vivo* suggests a link between Smo localization in primary cilia and rostral turning of dI1 commissural axons

To further support the implication of primary cilia in this pathway, we used our previously described *ex vivo* culture system to visualize dI1 axon behavior during navigation in real time (Fig. 5A) (Dumoulin et al., 2021). We first confirmed that dI1 neurons still carried a primary cilium after 1 day *ex vivo* in this culture system. Indeed, we could reveal the presence of primary cilia on dI1 neurons (expressing tdTomato-F) by staining for the ciliary marker Arl13b (white arrows, Fig. 5B).

**Figure 5.**
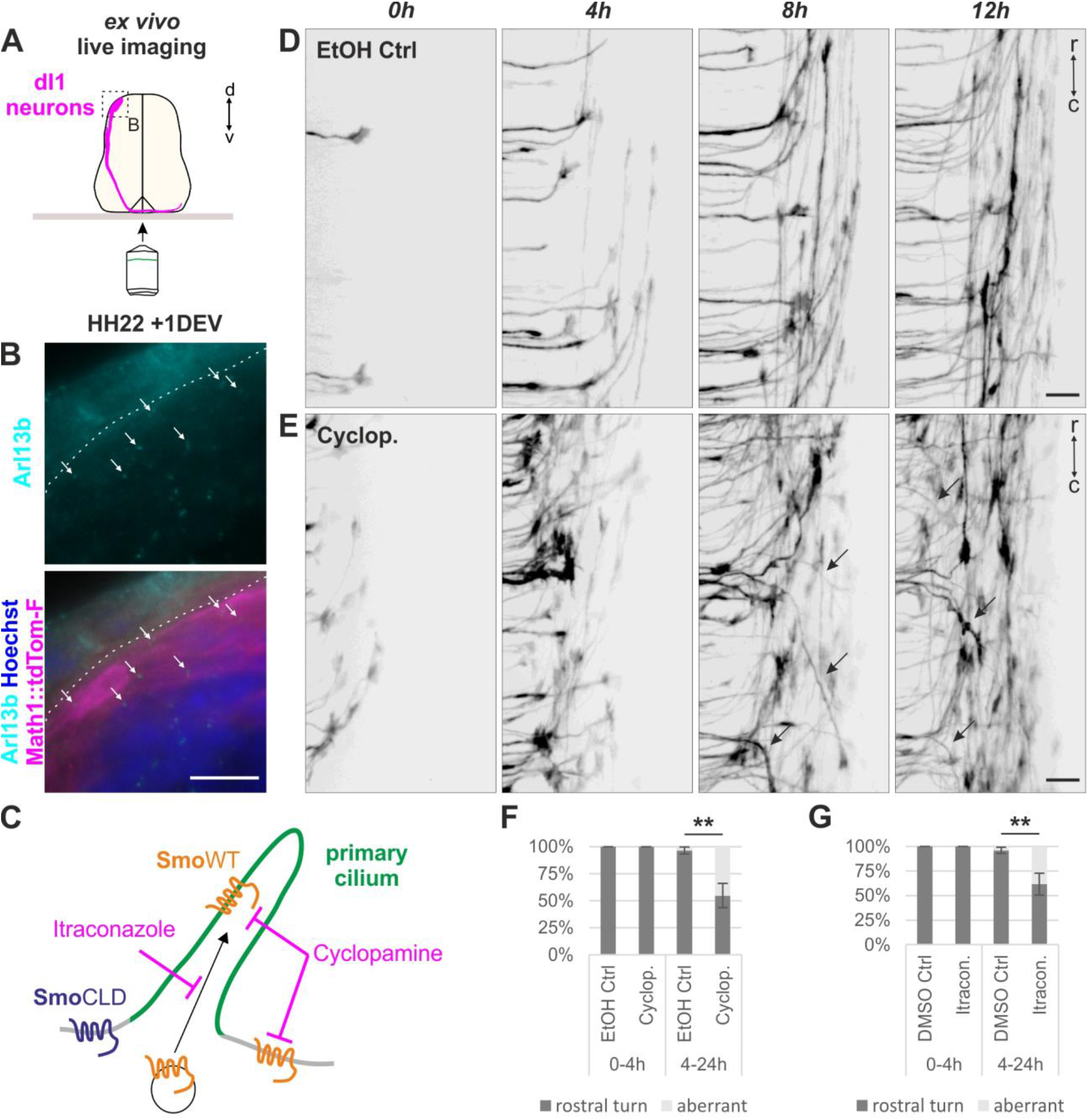
Ex vivo live imaging of dI1 commissural axons after Smo inhibition suggests a link between Smo localization in primary cilia and rostral turning of dI1 commissural axons. (A) Schematic depicting the ex vivo experimental set up for culture and visualization of dI1 axons guidance in the intact spinal cord. (B) After 24h of ex vivo culture, primary cilia were revealed by immunohistochemistry on transverse sections with the marker Arl13b (white arrows, cyan) on Math1-positive dI1 neuron cell bodies (magenta). Nuclei were counterstained with Hoechst (blue). (C) Schematic of ex vivo Smoothened (Smo) inhibition with the pharmacological blockers Cyclopamine (general inhibition of Smo) and Itraconazole (inhibition of Smo entry into the primary cilium). SmoCLD, a mutant Smo, cannot enter the primary cilium. (D-E) Snapshots of the live monitoring of dI1 commissural axons guidance at the contralateral floorplate border revealed aberrant guidance phenotypes in the presence of 15 μM Cyclopamine. Axons were turning caudally instead of rostrally (black arrows, E) compared to ethanol-treated vehicle control (D). (F-G) Quantification of the axon guidance phenotype at the single axon level at the contralateral floorplate border in the presence or absence of Cyclopamine or Itraconazole. No (0%) aberrant phenotype at the contra-lateral floorplate border was seen the first 4h of imaging in both Cyclopamine and Itraconazole-treated samples as well as in control conditions. However, a significant and similar increase of aberrant phenotypes for both inhibitors was found compared to controls between 4 and 24h of imaging (unpaired T-test). N(embryos)= 3 for each condition; n(axons)= 19 (EtOH Ctrl^0-4h^), 36 (Cyclopamine^0-4h^), 198 (EtOH Ctrl^4-24h^), 184 (Cyclopamine^4-24h^), 39 (DSMO Ctrl^0-4h^), 40 (Itraconazole^0-4h^), 142 (DSMO Ctrl^4-24h^) and 234 (Itraconazole^4-24h^). p<0.01 (**). DEV, day ex vivo; r, rostral; c, caudal; Cyclop., Cyclopamine; Itracon., Itraconazole. Scale bars: 10 μm (B) and 20 μm (D,E). Source data and statistics are available in Source Data spreadsheet.

Transcriptional output in primary cilia-dependent Shh signaling involves Smo translocation into the cilium (Briscoe and Thérond, 2013). Moreover, Smo was shown to be required for commissural axon guidance along the longitudinal axis (Parra and Zou, 2010; Yam et al., 2012). Thus, we took advantage of the *ex vivo* culture system and applied pharmacological blockers to inhibit Smo function during midline crossing (Fig. 5C). The presence of Cyclopamine in the culture medium blocked Smo function inside and outside the primary cilium (Fig. 5C). We could observe that dI1 commissural axons exiting the floorplate randomly turned into the longitudinal axis with a considerable number of axons turning caudally instead of rostrally (black arrows, Fig. 5E). These axons formed a disorganized post-crossing segment in comparison to the ethanol (EtOH)-treated control samples in which the large majority of axons turned rostrally (Fig. 5D, Movie S1). Remarkably, the Cyclopamine-mediated inhibition of Smo did not induce instantaneous aberrant guidance of commissural axons. There was no obvious effect on commissural axons exiting the floorplate during the first 4h of inhibition (Movie S1, Fig. 5E). In fact, dI1 growth cones that were traversing the floorplate, or were about to exit it, when the inhibitor was added all turned normally in rostral direction at the contralateral border (arrowheads, Movie S2). Mostly the ones that were not in the floorplate at the beginning of live imaging showed aberrant phenotypes (open arrowheads, Movie S2). Quantifications revealed that during the first 4 hours of time-lapse recording, there was no aberrant phenotype in both Cyclopamine- and control-treated samples (Fig. 5F, p>0.05). In contrast, between 4 and 24h of recording, the number of axons that turned caudally or stalled at the floorplate exit site was significantly higher (45 ± 11%) than in control (3 ± 3%, mean ± standard deviation, Fig. 5F, p<0.01). The fact that the Cyclopamine-mediated inhibition of Smo did not have an immediate effect on axon guidance was in agreement with our model that the aberrant phenotypes seen after at least 4 hours of culture (4.5 hours of inhibition) were caused by a transcriptional function of Smo. This also raised the possibility that these phenotypes were not caused by a growth cone-localized function of Smo as described for pre-crossing commissural axons (Yam et al., 2009). In line with our hypothesis, inhibition of Smo entry into the cilium with Itraconazole (Fig. 5C) (Kim et al., 2010) resulted in a very similar outcome. Axons behaved normally for at least the first 4h of culture with 100% of axons turning rostrally (Fig. 5G). After this time point, a significant number of axons (38 ± 8%) turned caudally or stalled at the contralateral floorplate border compared to the vehicle-treated control (4 ± 2%, mean ± standard deviation, DMSO, Movie S3, Fig. 5G, p<0.01).

Taken together, the real-time monitoring of axon guidance combined with pharmacological blockers of Smo suggest a function of Smo outside the growth cone but inside the primary cilium of dI1 neurons to allow correct commissural axon guidance at the post-crossing level.

### Smo localization in the primary cilium is required for the induction of Hhip transcription in dI1 neurons and proper axon guidance *in vivo*

We previously showed that the expression of a constitutively active Smo in the spinal cord up-regulated *Hhip* mRNA expression in dI1 neurons, suggesting a role for Smo in the Shh-Glypican-1-Hhip signaling pathway upstream of Hhip (Wilson and Stoeckli, 2013). To further decipher the role of Smo in this pathway, we investigated *Hhip* expression in embryos in which Smo localization to the cilium was perturbed. To achieve this, we knocked down endogenous chicken Smo, using artificial microRNA/shRNA (miSmo; Fig. S6), and then attempted to rescue *Hhip* expression by co-electroporating constructs encoding wildtype human Smo (hSmoWT), or a Smo mutant with two amino acid substitutions which cause defective ciliary localization (hSmoCLD). In contrast to hSmoWT, hSmoCLD cannot activate the transcription-dependent Shh response (Bijlsma et al., 2012; Corbit et al., 2005). Supporting the idea that *Hhip* induction in commissural neurons requires the cilium, we found that hSmoWT, but not hSmoCLD, could rescue *Hhip* expression in dI1 neurons following the knockdown of endogenous Smo on one side of the spinal cord (Fig. 6). We found that the average normalized ratio of *Hhip* expression in dI1 neurons after hSmoWT rescue was significantly higher than the one after either simple knockdown or rescue with hSmoCLD (Fig. 6D, p<0.05). Furthermore, expression of hSmoWT, but not hSmoCLD, rescued the axon guidance phenotypes induced by silencing endogenous *Smo* (Fig. 6E). After silencing *Smo* or expression of hSmoCLD after Smo knockdown, we found normal axon behavior at only 31 ± 20% and 25 ± 8% of the DiI injection sites (mean ± standard deviation). Only the rescue with hSmoWT brought the percentage of normal phenotype to a level similar to controls (see Fig. 3C) with 64 ± 17% of the DiI sites with normal axonal behavior at the ventral midline (mean ± standard deviation, Fig. 6E, p<0.01 versus miSmo and p<0.05 versus miSmo+hSmoCLD). Together, these results demonstrate that *Hhip* induction in the dorsal spinal cord relies on the ability of Smo to localize to the cilium. As *Hhip* mediates post-crossing commissural axon guidance (Bourikas et al., 2005; Wilson and Stoeckli, 2013), these results reveal a critical requirement for primary cilium-mediated Shh signaling in axonal navigation.

**Figure 6.**
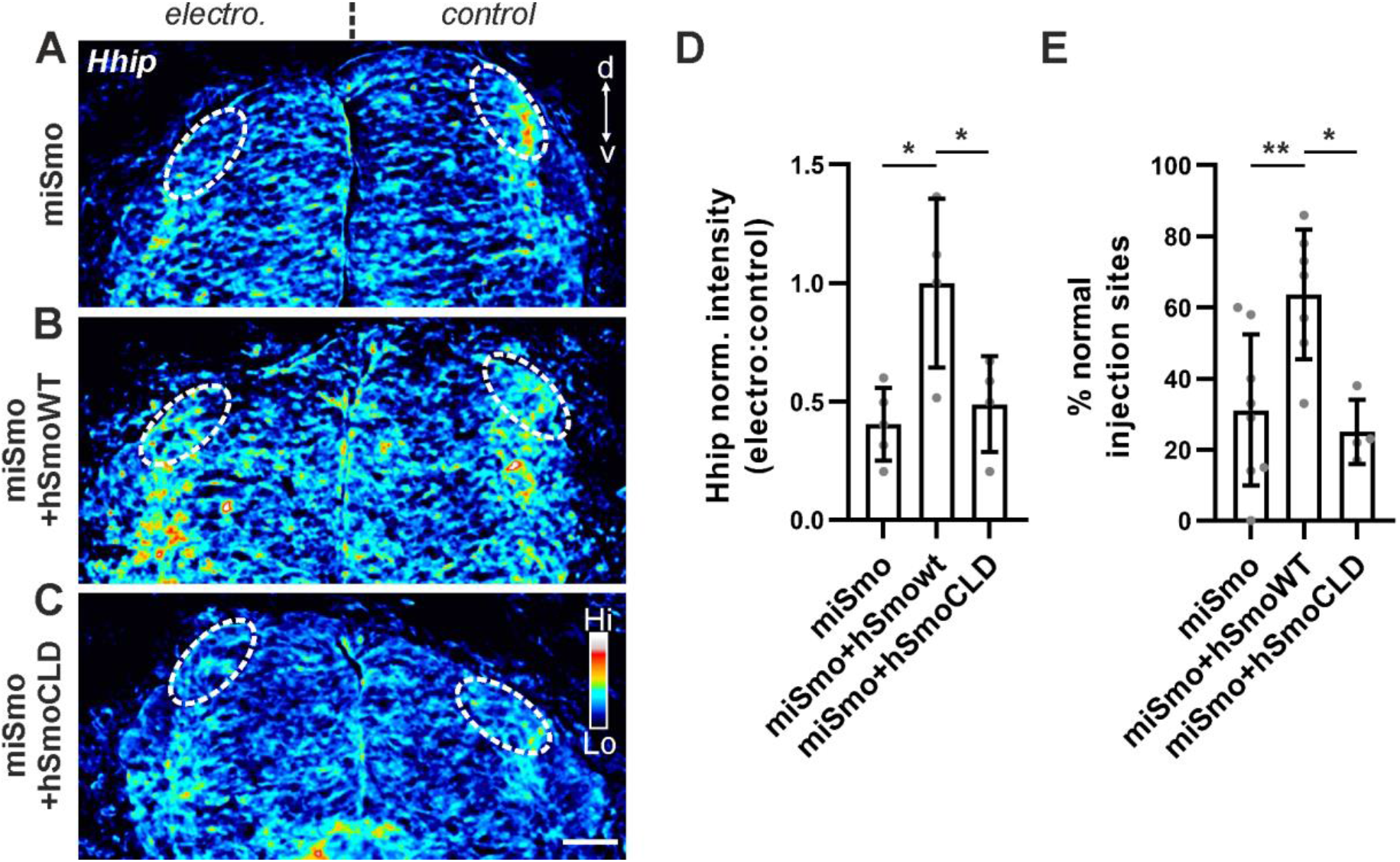
Smo localization in the primary cilium is required for transcription of Hhip in dI1 neurons and proper axon guidance in vivo. (A-C) Heat-map in situ hybridization pictures of HH25 dorsal spinal cords showing Hhip mRNA. Co-electroporation of wild-type human Smo (hSmoWT) rescued Hhip expression that was lost after Smo knockdown (A) in dI1 neurons (dashed ovals) compared to control side (B) whereas a SmoCLD mutant could not (C). (D) Quantification of Hhip in situ hybridization in dI1 neurons represented as a ratio between electroporated versus control side of the spinal cord and normalized to the average ratio of miSmo+hSmoWT condition. One-way ANOVA with Tukey’s multiple comparisons test. N(embryos)= 5 (miSmo), 4 (miSmo+hSmoWT) and 4 (miSmo+hSmoCLD). Note that each dot represents the average normalized ratio per embryo. (E) Rescue of Hhip expression after downregulation of Smo by co-electroporation of hSmoWT also rescued miSmo-induced axon guidance errors. Co-electroporation of cilia-localization-defective hSmoCLD was not able to rescue axon guidance. One-way ANOVA with Tukey’s multiple comparisons test. N(embryos) = 8 (miSmo), 7 (miSmo+hSmoWT) and 4 (miSmo+hSmoCLD). p<0.01 (**), p<0.05 (*) and p>0.05 (ns). d, dorsal; v, ventral; Hi, high; Lo, low; electro, electroporated. Scale bar: 50 μm. Source data and statistics are available in Source Data spreadsheet.

### Shh is transported from the axonal compartment to the soma of commissural neurons *in vitro*

During the time window of midline crossing by commissural axons, Shh is exclusively expressed in the floorplate (Bourikas et al., 2005). The requirement for the ciliary genes *Ift88 and Ift52*, as well as Smo localization to the primary cilium to trigger the Shh-Glypican-1-Hhip signaling cascade in dI1 neurons imply that a retrograde signal might come from the growth cones and travel to the soma, where the primary cilium is localized, when commissural axons are reaching the ventral midline of the CNS. This means that in order to trigger Smo translocation to the primary cilium, Shh needs to be transported from the growth cone to the soma, where it can bind to its receptor Patched, and thus, induce its ciliary exit.

To investigate this aspect we utilized microfluidic chambers with two chambers separated by microgrooves to culture dissociated commissural neurons dissected from HH25-26 chicken spinal cords. Neurons were seeded in the soma chamber and cultured for 8 days *in vitro* to allow their axons to cross the microgrooves and populate the axonal chamber (Fig. 7A). With this experimental setup, we could separate commissural axons from their soma. Importantly, after such a long time in culture, commissural neurons still carried a primary cilium as shown by co-staining of the primary cilium marker Arl13b and the neuronal marker neurofilament-M (arrows, Fig. 7B). To assess whether Shh could be transported from the axons to the soma of commissural neurons, we incubated the axons in the axonal chamber with recombinant ShhN fused to a 6xHis tag for 7 hours. Cultures were then washed and stained for Shh with an antibody against the His tag, with antibodies against Axonin-1 to label commissural axons or neurofilament-M (Fig. 7C).

**Figure 7.**
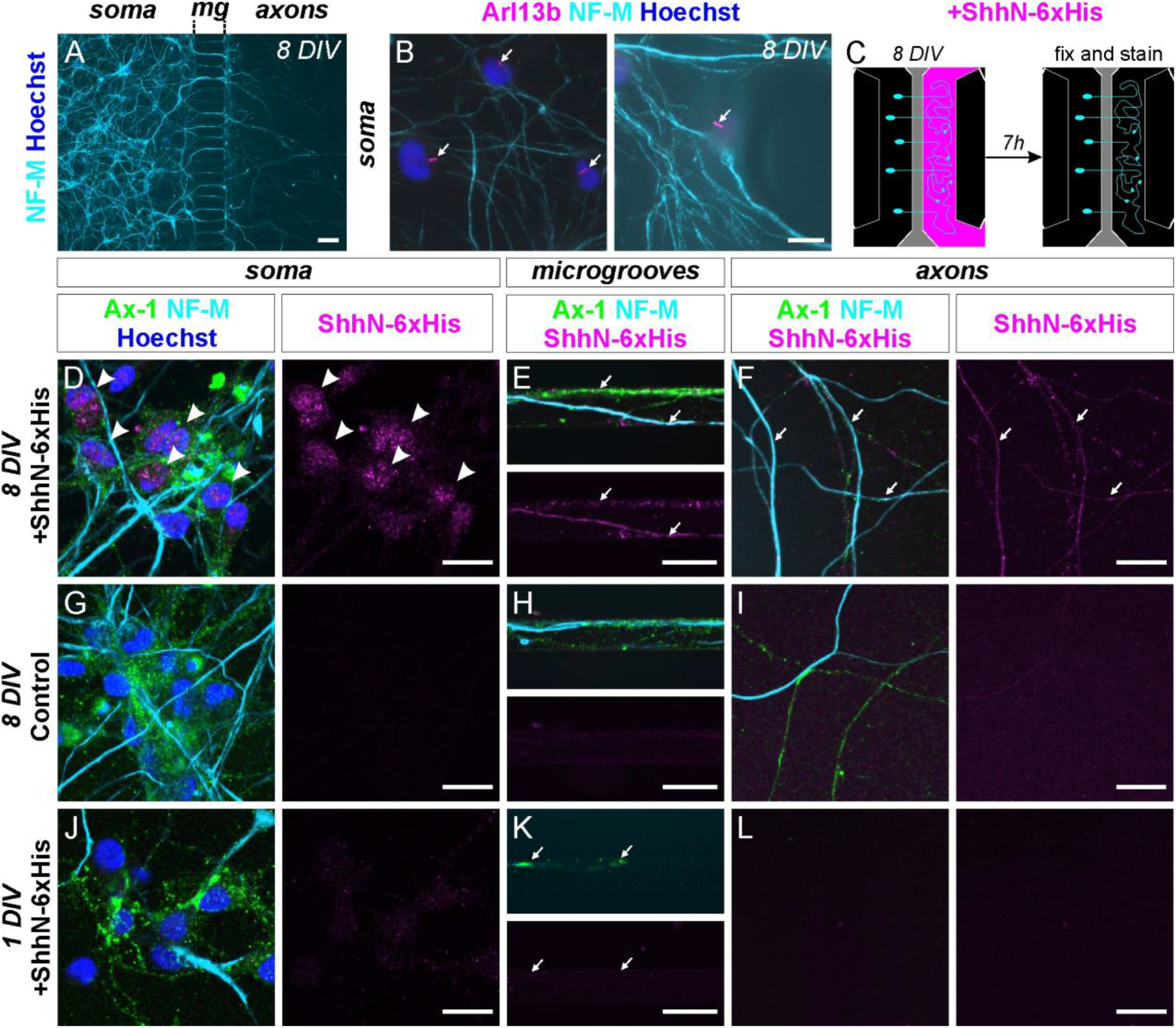
Shh is transported from the axonal compartment to the soma of commissural neurons cultured in microfluidic chambers. (A) Immunostaining of commissural neurons with neurofilament-M (cyan) cultured for 8 days in vitro in compartmentalized microfluidic chambers showing that axons could populate the axonal compartment. (B) Commissural neurons stained with neurofilament-M (cyan) still carried a primary cilium (white arrows), as shown by Arl13b staining (magenta). (C) Schematic of the experimental setup used to assess the localization of recombinant ShhN tagged with 6xHis (ShhN-6xHis, magenta). Commissural neurons (cyan) were cultured for 8 days in vitro before ShhN-6xHis was added for 7h in the axonal compartment. After that, neurons were washed, fixed and stained for neuronal markers neurofilament-M (cyan), commissural neuron marker Axonin-1 (green), and His tag (magenta), as shown in (D-J). (D) After 7h of incubation recombinant Shh (magenta) could be seen at the axonal level in the axonal compartment and microgrooves (white arrows) as well as at the soma level of commissural neurons (white arrowheads). (G) A first control was performed without application of the recombinant Shh in the axonal compartment leading to no His staining in all compartments, showing the specificity of the His antibody. (H) Most importantly, commissural neurons cultured for only one day in vitro that did not yet have the time to project their axons to the axonal compartment (white arrows point to growth cones traversing the microgrooves at the time of fixation) did not show any accumulation of recombinant Shh (magenta) at their somas (white arrowheads). This confirmed the proper fluidic compartmentalization between chambers. Nuclei were counterstained with Hoechst (blue). DIV, days in vitro; mg, microgrooves; Ax-1, Axonin-1; NF-M, neurofilament-M. Scale bars: 100 μm (A), 10 μm (B) and 20 μm (D-L).

We detected ShhN-6xHis at the axonal level in the axonal compartment and in the microgrooves between chambers (arrows, Fig. 7E,F) and also at the soma level of commissural neurons (arrowheads, Fig. 7D). We could not see any signal in control samples that were not incubated with recombinant Shh (Fig. 7G-I). Moreover, in microfluidic chambers containing commissural neurons that were cultured for only one day *in vitro* (Fig. 7J-L), and therefore did not have axons reaching the axonal chamber yet (arrows, Fig. 7K), we did not see any recombinant Shh at the soma level (arrowheads, Fig. 7J). This means that both soma and axonal chambers were fluidically isolated from each other during the 7-hour incubation time.

These results demonstrate that Shh can be retrogradely transported along axons of cultured commissural neurons, suggesting that Shh could reach the cell soma, when commissural axons are contacting the Shh-expressing floorplate *in vivo.* Taken together, our experiments demonstrate a primary cilium-dependent role of Shh in the induction of Hhip, its own receptor for the post-crossing phase of axon guidance.

## Discussion

Genes related to primary cilia formation, trafficking, and maintenance play crucial roles during development of the nervous system at many levels (Hasenpusch-Theil and Theil, 2021; Park et al., 2019; Suciu and Caspary, 2021). The use of animal models, especially the mouse, showed that mutation of ciliary genes led to aberrant formation of axonal tracts in the brain as well as axonal targeting in the CNS and PNS (Asadollahi et al., 2018; Green et al., 2018; Guo et al., 2019; Tadenev et al., 2011). The axonal tract malformations are reminiscent of the ones observed in patients with ciliopathies, such as Joubert syndrome (Sattar and Gleeson, 2011). Moreover, it was recently reported that loss of the ciliary genes Arl13b or Inpp5e had an impact on the development of callosal axons that cross the cortical midline to from the corpus callosum (Guo et al., 2019). This raised the possibility that these genes, and therefore a functional cilium, was involved in axonal navigation of intermediate targets in the CNS.

The developing spinal cord of mouse and chicken embryos enabled us to investigate the role played by the primary cilium in dI1 commissural neurons within the time window when their axons reached and crossed the intermediate target - the floorplate. We showed that these neurons carried a primary cilium when their axons were crossing the midline in both chicken and mouse embryos (Fig. 1). The use of a hypomorphic mouse model of the IFTB gene Ift88 with reduced ciliation in dI1 neurons revealed axon guidance defects at the exit of the CNS ventral midline (Fig. 2). Although these observations were compatible with a role for primary cilia in regulating the guidance of dI1 neurons at an intermediate target, the ventral patterning defects of the neural tube and loss of Shh in the floorplate in this mutant precluded any clear conclusion. Thanks to the spatiotemporal precision of *in ovo* RNAi-mediated knockdown of Ift88 in the dorsal chicken spinal cord, we could demonstrate a cell-autonomous role for Ift88 in the guidance of dI1 axons (Fig. 3). This showed that the loss of Ift88 had an impact on post-crossing axons which were mostly unable to turn rostrally (Fig. 3E,F). This phenotype was compatible with a loss of axonal sensitivity to gradients involved in their rostral turning, such as Shh (Bourikas et al., 2005). Ift88 reduction in the hypomorphic mouse model, or silencing of Ift88 at E3 in embryonic chicken dI1 neurons, did not have a detectable impact on pre-crossing dI1 axon guidance or growth rate, as reported in recent studies after manipulating Arl13b in primary cilia of interneurons at early stages in the chicken neural tube (Fig. 3 and S3), or in axons of deep cerebellar nuclei (Guo et al., 2019; Toro-Tapia and Das, 2020). Interestingly, Ift88 has been previously shown to be cell-autonomously required for the guidance of olfactory sensory axons projecting to dorsal glomeruli (Green et al., 2018). Note that the mild phenotype obtained after ventral knock-down of Ift88 might indicate a contribution of the primary cilium in maintaining the polarization of floorplate cells, as the apical end feet of these cells bear a primary cilium pointing towards the central canal (Fig. 3H,I) (Cruz et al., 2010).

Importantly, by investigating *Hhip* expression and function in dI1 neurons we could show that Ift88 functions upstream of Hhip, allowing its transient expression and engagement in axon guidance *in vivo* (Fig. 4). This also suggested the involvement of a functional primary cilium in long-range guidance of dI1 axons via Shh signaling (Wilson and Stoeckli, 2013). However, a cilium-independent role for Ift88 in this system cannot be excluded. In fact, the ciliary protein Arl13b has been recently reported to play a role outside the cilium in the guidance of pre-crossing dI1 axons in a transcription-independent manner in the mouse (Ferent et al., 2019). Both the transcriptional change in *Hhip* expression after Ift88 knockdown and Smo localization into the cilium, which is required for both *Hhip* mRNA expression in dI1 neurons and correct dI1 axon guidance *ex vivo* and *in vivo* (Fig. 4, 5, and 6), strongly support a model in which the primary cilium functions upstream of *Hhip* in dI1 neurons. Thus, our data support a role for the primary cilium in switching commissural dI1 axons’ responsiveness to Shh from attraction to repulsion at the intermediate target (Fig. 8). This implies a switch of Shh signaling in these neurons from a transcription-independent to a transcription-dependent manner (Fig. 8) (Yam et al., 2012). The average time dI1 axons take from their first contact with the floorplate (Shh producing cells) to initiating the rostral turning at the contralateral border (~7h) is compatible with a transcriptional switch of Shh receptors (Dumoulin et al., 2021). This also raises the question of which transcription factor(s) is/are required downstream of the primary cilium to induce *Hhip* transcription. In line with a transcription-dependent role for the primary cilium in this system, previous results supported that *Hhip* transcription induction in dI1 neurons might be Gli-dependent as Gli1 or 2 overexpression induced up-regulation of *Hhip* in the dorsal spinal cord (Wilson and Stoeckli, 2013). Moreover, a Hedgehog-insensitive dominant repressor of Smo abolished Hhip expression in dI1 neurons, further supporting transcription-dependent Shh signaling (Wilson and Stoeckli, 2013). Nevertheless, further experiments are required to assess in more detail whether the Gli-dependent transcriptional pathway is activated, when dI1 axons are crossing the floorplate, and whether this depends on a functional primary cilium.

**Figure 8.**
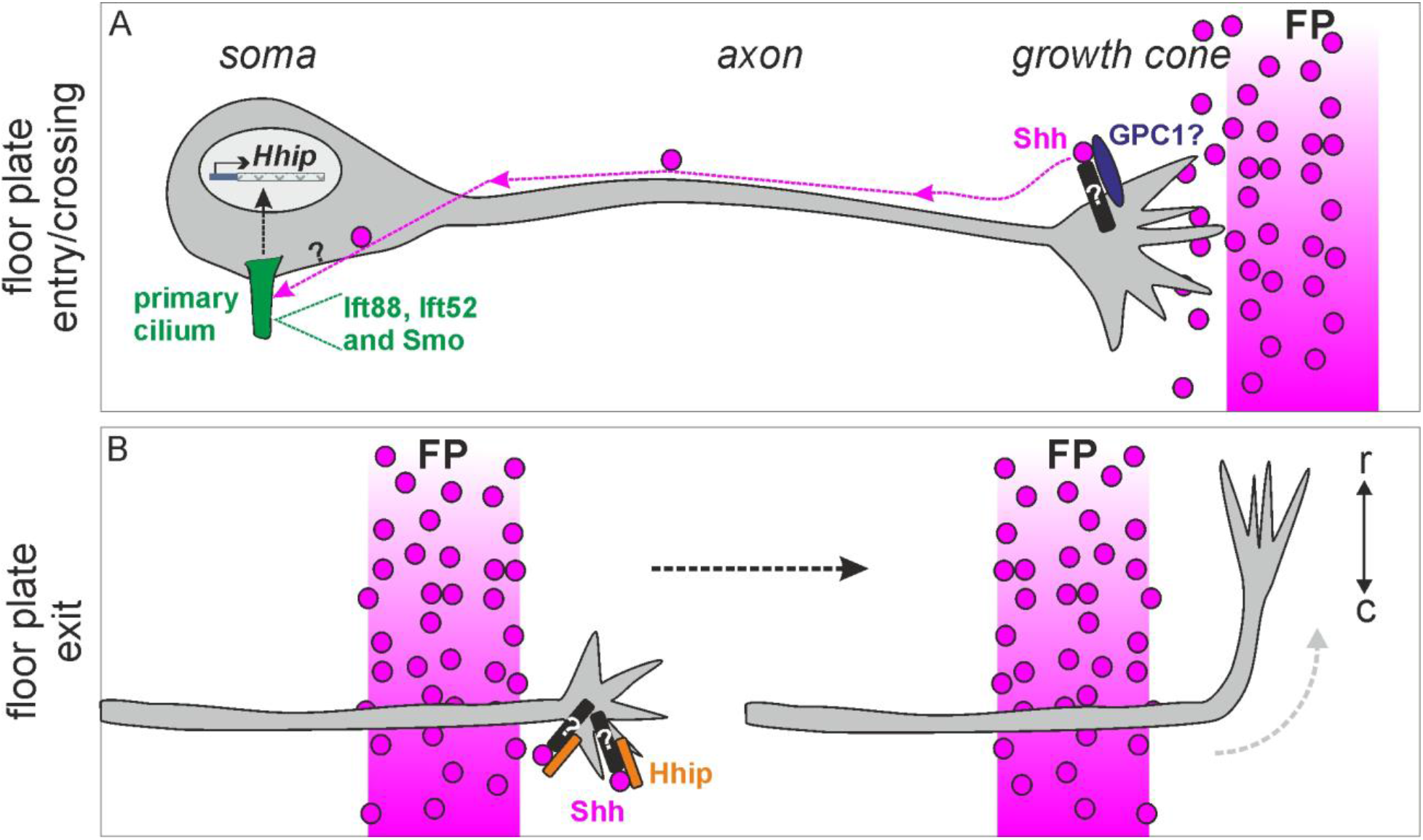
Model for the primary cilium’s function in Shh-mediated transcriptional regulation of Hhip and thus, the switch in responsiveness during long-range commissural axon guidance. (A) Our data support a signaling pathway in which Shh produced by the floorplate is retrogradely transported to the soma of commissural neurons, when their growth cone contacts the intermediate target. As the Glypican-1--Shh interaction was reported to be required for the induction of Hhip transcription, it is likely that Glypican-1 interacts with Shh at the growth cone and that by a yet unknown mechanism Shh is transported retrogradely in- or outside the axons. Ciliary proteins Ift88 and Ift52, and Smo localization to the cilium are required for correct commissural axon guidance. Ift88 as well as Smo in the cilium are both required for the induction of Hhip transcription. Together, this strongly supports a model in which the cilium plays a pivotal intermediate role in this signaling cascade. When Shh reaches the soma, it would induce Patched exit of the primary cilium, and in turn, cause Smo entry, leading to a transcriptional output. (B) Once the commissural axons exit the floorplate at the contralateral side, Hhip protein has to be expressed and transported to the growth cone surface, causing the repulsive response to the caudal^high^-rostral^low^ Shh gradient and the turn towards the brain. Black rods represent unknown co-receptors. FP, floorplate; GPC1, Glypican-1; r, rostral; c, caudal.

Our results also bring some new insights in how Shh might signal in a long-range manner in neurons. When dI1 axons reach the floorplate – the only source of Shh in the spinal cord at that time – their cilium is located around 400-500 μm away from it. We know that the Glypican1-Shh interaction is required for *Hhip* induction in dI1 neurons (Wilson and Stoeckli, 2013) and this study suggests that a functional primary cilium is required for *Hhip* induction. This implies that the Shh protein has to be transported to the cilium on the neuronal cell body. Indeed our microfluidic assays supported this hypothesis (Fig. 7).

Recently, Shh has been shown to be anterogradely transported by retinal ganglion cells via their axons and to be secreted in the optic chiasm, where it could act as a repellent for ipsilateral populations of retinal ganglion cell axons (Peng et al., 2018). Here, our results show that the dI1 population of commissural neurons is able to retrieve and transport Shh for a long distance in a retrograde manner. Future experiments will be required to assess whether Shh is transported at the surface of dI1 axons or whether it is internalized. By analogy to other systems, we could imagine two plausible scenarios: (1) Shh could be internalized at the growth cone, transported internally, and secreted at the soma, as it is the case in Shh transcytosis in epithelial cells (Ho and Stearns, 2021). (2) as an interaction between Glypican 1 and Shh is required in our system, we can imagine that Shh is transported retrogradely at the surface of dI1 axons, as it is the case in Drosophila with the Glypican orthologue Dally-like protein (dlp), which is present in cytonemes of receiving cells (long signaling filopodia) (González-Méndez et al., 2017). In our case, dI1 axons might play the role of a receiving cytoneme for long-range transport of Shh to the neuronal soma (Fig. 8A).

Taken together, we propose a signaling model in which Shh, secreted by the floorplate, is retrogradely transported by dI1 axons after contact with the intermediate target. Thus, Shh reaches their soma and transmits a signal at the primary cilium, which will induce transcriptional activation of *Hhip* (Fig. 8A). At the time when dI1 axons exit the floorplate, Hhip protein is expressed and can act as a Shh receptor on the growth cone surface to respond to the repulsive Shh gradient and make the axon turn rostrally towards the brain (Fig. 8B) (Bourikas et al., 2005). These results shed light on a molecular mechanism of axon guidance that might contribute to our understanding of the etiology of human ciliopathies. The results presented here are in agreement with our recent findings on the role of C5ORF42 (also termed CPLANE1 or JBTS17) in neural circuit formation (Asadollahi et al., 2018). Patients with mutations in this gene were diagnosed with a subtype of Joubert syndrome, OFDVI (oro-facial-digital syndrome type VI) (Bayram et al., 2015; Romani et al., 2015). Fibroblasts derived from patients exhibited a lack of cilia. Furthermore, the phenotypes of the patients with a mutation in C5ORF42 were very similar to those with a mutation in Kif7, another ciliary gene. Most importantly, the axon guidance phenotypes observed in chicken embryos after silencing C5ORF42 were very similar to those described here for embryos lacking functional cilia in the absence of Ift88 (Asadollahi et al., 2018). Taken together, these studies suggest that primary cilium-mediated, transcription-dependent Shh signaling is required for neural circuit formation.

## Materials and Methods

### Animals

All experiments were approved and carried out according to the guidelines of the Cantonal Veterinary Office Zurich. Cobblestone (*cbs*) mice were generated as described (Willaredt et al., 2008) and genotyped by PCR, using genomic DNA. The following primer sets were used: forward1: 5’-TTGACATCTGGATATGACAATGC, reverse1: 5’-TGTGCATGTTTGTGTACATATGTG; and forward2: 5’-TGGTGTCTCCTTCGGAATTT, reverse2: 5’-TAAATGTAAAAGGTAAAGGCAATGG. Noon of the day of the vaginal plug was designated embryonic day 0.5 (E0.5). Fertilized chicken eggs were obtained from a local farm and staged according to Hamburger and Hamilton (Hamburger and Hamilton, 1951).

### Assessment of the longitudinal Shh gradient

Wildtype E12.5 NMRI mice were sacrificed, pinned flat in the supine position, and fixed in 4% paraformaldehyde (PFA) in PBS for 2 hours at room temperature, before rinsing in phosphate buffered saline (PBS), and soaking in 25% sucrose in PBS overnight. Embryos were mounted and frozen in TissueTek O.C.T. compound. Twenty-five μm-thick transverse sections were collected, with 10 sections on each slide, at 400 μm intervals. The collected area spanned from the hindlimbs to the forelimbs. After *in situ* hybridization, *Shh* intensity in the floorplate was calculated using ImageJ software (NIH, USA). *In situ* images were inverted, the floorplate area was selected, and the Mean Grey Value in the selected area was measured. The selected border was moved to an area of the section in the dorsal spinal cord (where no *Shh* is present) and the Mean Gray Value was resampled as a background measurement. This background value was subtracted from the floorplate measurement to give a final *Shh* intensity measurement in each section. *Shh* measurements along the longitudinal axis were normalized within each embryo by dividing the measured intensity at the mid-trunk and forelimb levels by the value obtained in the hindlimb region. Thus, normalized Shh intensity values of <1 in the mid-trunk and forelimb levels indicated weaker expression than in the hindlimb region of the same embryo. Twelve embryos were used in the analysis. Data is presented as mean ± SEM and was subjected to a single-sample T-test against an intensity of 1 (arbitrary units).

### Long double-stranded RNA and siRNA

Chicken ESTs of *Ift88* (ChEST 972d21) and *Ift52* (ChEST 490p6) were obtained from Source BioScience LifeSciences (Nottingham, UK). The *Ift88* plasmid was linearized with EcoRI or NotI, while the *Ift52* plasmid was linearized with KpnI or SacI (New England Biolabs), for the synthesis of sense and antisense single-stranded RNAs (ssRNA) with T7 and T3 RNA polymerase (Promega), respectively. Equal amounts of purified ssRNAs were annealed by allowing the solution to cool slowly to room temperature after heating at 95°C for 10 minutes. Successful double-strand formation was verified by gel electrophoresis. As a control, we used dsRNA synthesized from a 209 bp fragment of red fluorescent protein (mRFP1). Long dsRNAs were electroporated at a concentration of 400 ng/μl in PBS, together with 40 ng/μl of a transfection reporter plasmid encoding humanized *Renilla* GFP (hrGFPII; Stratagene) and 0.01% Fast Green (AppliChem).

We used an *in vitro* reporter approach to test the effectiveness of target gene knockdown of the different dsRNA sequences (Fig. S5 and S6). The *Ift88* or *Ift52* EST sequences were cloned into a CMV-driven vector, between the stop codon of *EGFP* and a poly-A tail. Long dsRNAs were digested to siRNAs and purified according to the instructions of ShortCut^®^ RNase III (NEB). We then co-transfected COS7 cells with the Ift reporter constructs, together with EBFP2 (as a transfection control) and siRNA against *RFP*, *Ift52* or *Ift88*, using Lipofectamine 2000 (Invitrogen) in LabTekII chamber slides (Nunc). Expression levels of EBFP2 and EGFP were assessed 24 hours later. Ten images in each condition were quantified in ImageJ (Mean Gray Values), then normalized to transfection levels (EGFP:EBFP2) and subjected to statistical analyses (One-way ANOVA with Tukey’s multiple comparisons test).

### Artificial miRNAs

Plasmids encoding artificial miRNAs against chicken *Smoothened* (Smo) and Luciferase were synthesized as described (Wilson and Stoeckli, 2011). The target sequences were: (miSmo) 5’-AAGTGCAGAACATCAAGTTCA; (miLuc, against firefly Luciferase; (Wilson and Stoeckli, 2011)) 5’-CGTGGATTACGTCGCCAGTCAA. For *in ovo* electroporations, we used plasmids (250 ng/μl) containing a β-actin promoter driving the expression of hrGFPII and an artifical miRNA, followed by a poly-A tail mix in PBS and Fast Green as described above.

The effectiveness of the artificial miRNAs was tested indirectly using a similar method to that described above for the dsRNAs (Fig. S6; (Wilson and Stoeckli, 2011)). miRNAs were cloned into pRFPRNAiC vectors and co-transfected into COS7 cells along with *Smo* reporter constructs, in which a 2.2 kb fragment of chick *Smo* was cloned downstream of EGFP in either the 5’-3’ direction or, as a negative control, in the opposite orientation. Expression levels of RFP and EGFP were assessed 24 hours later. Fifteen images in each condition were quantified in ImageJ (Mean Gray Values), normalized to transfection levels (EGFP:RFP) and subjected to statistical analyses (unpaired T-test).

### SmoWT and SmoCLD constructs

A cDNA clone containing human *Smo-M2*, a constitutively active form of human *Smo* with a single point mutation (W535L) was kindly provided by J. Briscoe. This construct was tagged with the first 55 amino acids of human herpes virus glycoprotein D (HHV gD1) at the N-terminus, thus all subsequent hSmo constructs were detectable using an anti-gD1 antibody. *hSmo-M2* was mutagenized to *hSmoWT* using the primers 5’-CCATGAGCACCTGGGTCTGGACCAAG and 5’-CTTGGTCCAGACCCAGGTGCTCATGG. To make *hSmoCLD*, we subsequently mutagenized two sequential amino acids, W545A and R546A, using the primers 5’-CATCGCGGCGCGTACCTGGTGCAG and 5’-GTACGCGCCGCGATGAGCAGCGTG. These amino acids were equivalent to W549A, R550A mutations in mouse *Smo*, which cause a ciliary localization defect (Bijlsma et al., 2012; Corbit et al., 2005). We confirmed *in vitro* that the *hSmo* constructs were not effectively downregulated by miSmo, which was designed against chicken *Smo* (Fig. S6D). For *in ovo* electroporations (using 50 ng/μl), the *Smo* variants were cloned into pMES, which is driven by a β-actin promoter and contains an IRES-GFP sequence to identify the transfected cells. The *Smo* variants were excised from pRK7 using HindIII (blunted) and EcoRI, while pMES was digested with XbaI (blunted) and EcoRI. The fragments were ligated using T4 DNA ligase (all enzymes from NEB).

### Electroporation and assessment of axon guidance phenotypes

A detailed video protocol demonstrating the electroporation, dissection and DiI injection steps in chicken embryos is available online: http://www.jove.com/video/4384 (Wilson and Stoeckli, 2012). In brief, embryos were injected and electroporated at HH17-18, using a BTX ECM830 square-wave electroporator (5 pulses of 25 V, 50 msec duration, 1 sec interpulse interval). Targeted dorsal and ventral electroporations were achieved by careful positioning of the electrodes relative to the neural tube, and successful targeting was verified by hrGFPII expression. The resulting axon guidance phenotypes were assessed by axonal tracing with DiI in open-book preparations of spinal cords. Note that the downregulation and rescues experiments of Smo were performed around HH15-16 to efficiently downregulate Smo. The spinal cords of E12.5 mice or HH25-26 chicken embryos were dissected, and ‘open-book’ preparations were made by cutting along the roof plate and pinning the spinal cord open with the basal sides down, as depicted in Fig. 2F. At least 7 embryos were examined in each condition by a person blind to the experimental condition. Fast-DiI (5 mg/ml in ethanol; Molecular Probes) was applied by focal injection into dorsal commissural neurons. Labeled axons at the midline were documented by confocal microscopy (Olympus DSU coupled to BX61 microscope). Only DiI injections sites that were in the appropriate location in the dorsal-most part of the spinal cord, and (for the chicken embryos) within the region expressing fluorescent protein, were included in the analysis. As it was impossible to count axons at individual injection sites, the percentage of axons displaying abnormalities was estimated, and the injection site was classified as showing a ‘FP stalling’ phenotype, if >50% of axons stalled within the floorplate, or a ‘post-crossing’ phenotype, if >50% of axons that reached the contralateral floorplate border failed to make a correct turning decision into the longitudinal axis. At a single abnormal DiI injection site, it was possible that more than one class of phenotypic error was observed. The total number of DiI sites in each condition was pooled and the percentage of normal injections sites were statistically compared across conditions.

### In situ hybridization

*In situ* hybridization and immunolabeling were performed as described (Mauti et al., 2006; Wilson and Stoeckli, 2011). All sense and antisense ISH probes were generated using SP6, T7 or T3 RNA polymerase, and DIG RNA Labeling mix (Roche). A plasmid containing a fragment of mouse cDNA for Shh was a gift from T. Thier. The chick *Hhip* probe was previously described (Bourikas et al., 2005). Images were acquired with a BX63 upright microscope (Olympus) and a 20x air objective (ACHN P 20x / 0.4, Olympus) and an Orca-R^2^ camera (Hamamatsu) with the Olympus CellSens Dimension 2.2 software.

For comparison of the expression patterns of *Shh* along the longitudinal axis of the spinal cords of WT mice, we embedded several fixed embryos in the same block, to enable a comparison of different embryos at the same axial level whilst minimizing slide-to-slide variability. The embryos were laid side-by-side in the supine position in O.C.T. compound (Tissue-Tek), and we aligned their hindlimbs and forelimbs perpendicular to the cutting surface before freezing. Twenty-five μm-thick cryostat sections at 400 μm intervals were collected on slides, such that each slide contained 10 sections that spanned from the hindlimb to the forelimb level. Analysis of relative *Shh* mRNA levels on the control and electroporated sides of the spinal cords was performed as described (Wilson and Stoeckli, 2013).

For *Hhip in situ* quantification in dI1 neurons of the electroporated versus the non-electroporated side on cryosections (Fig. 4 and 6), images were inverted and the mean value within a circle of 50-μm diameter positioned dorsally to the dorsal funiculus was measured on both control and electroporated side. Another value with the same circle was taken in the motor column of the un-electroporated (control) side. As *Hhip* is not expressed in the motor column (Wilson and Stoeckli, 2013) and as the density of cells in this area is quite similar to the area with the dI1 neurons, we used it as a measure of background. This background value was subtracted from each *Hhip* mean value in dI1 neurons and the ratio electroporated:control side was then calculated for each section. Note that sections with a mean value of *Hhip* in dI1 neurons inferior to the background value in the motor column were not taken into account. A minimum of 10 sections were quantified per embryo and the average ratio for each condition was normalized to the average ratio of GFP controls (Fig. 4C) or hSmoWT rescue (Fig. 6D). For comprehensive representation *in situ* images were inverted and Royal LUT implemented in ImageJ to highlight accumulation or reduction of mRNA in dI1 neurons (Fig. 4A,B and 6A-C).

### Immunohistochemistry for neuronal patterning

Mouse or chicken embryos were sacrificed, dissected and fixed in 4% PFA in PBS for 1h at room temperature, as described. After being washed 3 times 5 min in PBS, they were cryopreserved for 24h in 25% sucrose in PBS at 4°C and then embedded in O.C.T. compound (Tissue-Tek). From the trunk, 25-μm thick transverse sections were cut using a Cryostat (Leica). Sections were permeabilized for 10 min at room temperature with 0.1% Triton X-100 in PBS, blocked for 1h in 0.1% Triton X100, 5% FCS in PBS (blocking buffer). Cryosections were incubated overnight in the following primary antibodies diluted in blocking buffer: mouse anti-Pax3 (RRID: AB_528426), mouse anti-Islet1 (40.2D6, RRID: AB_528315), mouse anti-Shh (5E1, RRID: AB_2188307), mouse anti-HNF3β (4C7, RRID: AB_2278498), mouse anti-Lhx2 (PCRP-LHX2-1C11, RRID: AB_2618817), mouse anti-Nkx2.2 (74.5A5, RRID: AB_531794; all from Developmental Studies Hybridoma Bank), goat anti-GFP-FITC (1:400, Rockland, RRID: AB_218187) or rabbit anti-Axonin1 (1:1000, (Stoeckli and Landmesser, 1995)). The next day, sections were washed 3 times 10 min in 0.1% Triton X-100 in PBS at room temperature and then incubated for 2h with secondary antibodies diluted in blocking buffer: donkey anti-rabbit-IgG-Cy3 antibody (1:1000, Jackson ImmunoResearch, RRID:, AB_2307443), donkey anti-mouse-IgG-Cy5 antibody (1:1000, Jackson ImmunoResearch, RRID:AB_2338713) or donkey anti-mouse-IgG-Cy3 (1:1000, Jackson ImmunoResearch, RRID: AB_2340813). Finally, sections were washed 3 times 10 min in 0.1% Triton X-100 in PBS and 2 times 5 min in PBS before being mounted under a coverslip in Mowiol/DABCO. Images were acquired with a BX63 upright microscope (Olympus) and a 20x air objective (ACHN P 20x / 0.4, Olympus) and an Orca-R^2^ camera (Hamamatsu) with the Olympus CellSens Dimension 2.2 software.

### Immunohistochemistry of primary cilia

For staining of neuronal cilia in the neural tube, we had to modify certain steps of the protocol to increase signal to background of ciliary staining. Chicken embryos were sacrificed and dissected as described above in warm (37°C) PBS and then fixed at room temperature in pre-warmed 4% PFA in PBS for different times according to the stage (HH20, 35 min; HH22, 40 min; HH24, 40 min; HH26, 45 min). Note that this initial change in the protocol was not required for mouse embryos. The embedding and cryosections were performed as above as well as the immunostaining of dI1 neurons with Lhx2 or GFP. Once this staining was finished and sections washed with PBS, they were incubated for 2h in primary antibodies diluted in 5% FCS in PBS at room temperature to stain primary cilia with either rabbit anti-Arl13b (1:500, Proteintech, RRID:AB_2060867) or rabbit anti-Adenylyl cyclase III (ACIII, 1:500, Santa Cruz, RRID:AB_630839). Sections were then washed 3 times for 10 min in PBS and stained for 2h with donkey anti-rabbit-IgG-Cy3 antibody (1:1000, Jackson ImmunoResearch, RRID:, AB_2307443) and 2.5 μg/ml of Hoechst (Cat# H3570, Invitrogen) diluted in PBS. Finally, they were washed 3 times for 10 min in PBS and mounted as above. Images of cilia stainings were acquired with a BX61 upright microscope (Olympus) equipped with a spinning disk unit either with a 10x air objective (overviews; UPLFL PH 10x / 0.30, Olympus) or a 60x oil objective (PLAPON O 60x / 1.42, Olympus) and an Orca-R^2^ camera (Hamamatsu) with the Olympus CellSens Dimension 2.2 software.

### Live imaging of intact spinal cords

Live imaging of intact chicken spinal cords was performed as previously described (Dumoulin et al., 2021). In brief, the neural tube of HH17-18 embryos was injected and unilaterally electroporated *in ovo* as described above (25 volts, 5 × 50 ms pulses with 1-s interval between pulses) with 700 ng/μl Math1::tdTomato-F plasmid to label dI1 neurons and 30 ng/μl of β-actin::EGFP-F plasmid as control in all transfected cells (Dumoulin et al., 2021). Intact spinal cords were dissected at HH22 and cultured in a 100-μl drop of low-melting agarose with the ventral midline facing down on a 35-mm Ibidi μ-dish with glass bottom (Ibidi, Cat#81158) and spinal cord medium (MEM with Glutamax [Gibco] supplemented with 4 mg/ml Albumax [Gibco], 1 mM pyruvate [Sigma], 100 units/ml Penicillin and 100 μg/ml Streptomycin [Gibco]), as previously described (Dumoulin et al., 2021).

Intact spinal cords were cultured at 37°C, with 5% CO2 and 95% air in a PeCon cell vivo chamber (PeCon). CO2 percentage and temperature were controlled by the cell vivo CO2 controller and the temperature controller units, respectively (PeCon). Spinal cords were incubated for 30 minutes in the chamber before the live imaging acquisition was initiated with an IX83 inverted microscope equipped with a spinning disk unit (CSU-X1 10,000 rpm, Yokogawa) and a 20x air objective (UPLSAPO ×20/0.75, Olympus) and an Orca-Flash 4.0 camera (Hamamatsu) with the Olympus CellSens Dimension 2.2 software. We acquired one stack of 30-45 slices with 1.5-μm spacing every 15 min for 24 h with 488 nm and 561 nm emission channels, as well as the bright field channel to visualize the structure of the midline area. For the Smo inhibition experiments, either 15 μM of Cyclopamine diluted in ethanol (MedChemExpress, Cat# HY-17024) or 20 μM of Itraconazole diluted in DMSO (Sigma, Cat# I6657) as a final concentration was added to the spinal cord medium (1:000 dilution). Control conditions consisted of spinal cords cultured in 1:000 dilution of either ethanol (Cyclopamine control) or DMSO (Itraconazole control). As no obvious defects in midline crossing were detected upon Smo inhibition, we focused our quantification of aberrant phenotypes on the post-crossing segment of Math1-positive dI1 axons. An axon was considered to have an aberrant phenotype, if it stalled or turned caudally instead of rostrally, at the contra-lateral border of the floorplate. Tracing/counting of dI1 axons, processing of time-lapse video and video montages were performed with Fiji/ImageJ (Schindelin et al., 2012).

### Commissural neuron cultures in microfluidic chambers

The most dorsal part of the spinal cord, containing mostly dI1 commissural neurons, was cut from 6-7 open-book preparations of wild-type HH25-26 embryos in cold sterile PBS as previously described (Yusifov et al., 2021). Cells were dissociated with 0.25% Trypsin in PBS (Invitrogen, cat# 15090-046), containing 0.2% DNase (Roche, cat# 101 041 590 01), at 37°C for 20 min followed by a sequence of trituration in different pre-warmed media (37°C; MEM (Gibco) with 5% FCS (Gibco), MEM only, and finally in commissural neuron medium) with fire-polished Pasteur pipettes and centrifugation at 1000 rpm for 5 min at room temperature in-between. Commissural neuron medium contained MEM/Glutamax (Gibco, Cat# 41090-028) supplemented with 4 mg/ml Albumax (Gibco), N3 (100 μg/ml transferrin, 10 μg/ml insulin, 20 ng/ml triiodothyronine, 40 nM progesterone, 200 ng/ml corticosterone, 200 μM putrescine, 60 nM sodium selenite; all from Sigma) and 1 mM pyruvate (Sigma). Dissociated cells (130’000) in 20 μl commissural neuron medium volume were given in the upper left well of Xona chip 150 μm microfluidic chambers connected to the soma chamber (Xona, Cat# XC150). After 5 min at room temperature, 100 μl of medium were given to all wells. Plates were incubated at 37°C with 5% CO_2_ for 1 or 8 day. Every two days, 50% of the volume was changed in each well with freshly prepared medium. The microfluidic chips were coated with 20 μg/ml poly-L-lysine (Sigma, Cat# P-12374) and 20 μg/ml laminin (Invitrogen, Cat# 23017-015) following the protocol of the manufacturer (XonaChip™protocol for primary murine neurons). On the day of stimulation (1 or 8 day *in vitro*), culture medium was discarded from all wells, and 140 μl commissural medium were added to both left wells (connected to the soma chamber) and 130 μl of the same medium containing 2.5 μg/ml of recombinant ShhN-6xHis (R&D, Cat# 1845-SH; 1:40 dilution in medium of stock solution solubilized in 0.1% BSA) were given to both wells connected to the axonal chamber. Note that in control experiments without recombinant Shh (Fig. 7G-I) the equivalent amount of 0.1% BSA (1:40 dilution in medium) was given to the well connected to the axonal chamber. Cells were then incubated at 37°C with 5% CO2 for 7 h. Afterwards, the medium from the right side (axonal chamber) was discarded prior to the one in the soma chamber and then cells were washed once with pre-warmed (37°C) commissural neuron medium with first adding 100 μl in the left wells and 80 μl in the right wells. Then, cells were fixed with pre-warmed (37°C) 2% PFA in PBS and incubated at 37°C with 5% CO2 for 5 min prior to 15 min under same condition in 4% pre-warmed PFA in PBS. Finally, cells were washed 3 times for 5 min with PBS (150 μl per well) at room temperature and stored at 4°C until immunocytochemistry was performed.

### Immunocytochemistry microfluidic chambers

Importantly, for successful staining in microfluidic chamber microgrooves, the addition of any buffer in the wells was always 40 μl higher on one side of the dish (typically 150 μl in left wells and 130 μl in right wells). Neurons were first permeabilized with 0.1% Triton X-100 in PBS for 5 min at room temperature and washed 3 times 5 min with PBS. They were then blocked for 15 min in 5% FCS in PBS (blocking buffer). Samples were stained overnight with the following primary antibodies diluted in blocking buffer: mouse anti-neurofilament-M (1:1500, RMO270, Invitrogen, RRID:AB_2315286); rabbit anti-6xHisTag (1:2000, Rockland, Cat# 600-401-382) and goat anti-Axonin-1 (1:1000, (Stoeckli and Landmesser, 1995)). For Arl13b staining, samples were incubated only for 1h at room temperature with the primary antibody (rabbit anti-Arl13b, 1:1000, ProteinTech, 13967-1-AP, RRID:AB_2121979). After being washed 3 times 5 min with PBS at room temperature, neurons were stained 2 h (1h for Arl13b staining) at room temperature with secondary antibodies (same as used for immunohistochemistry) diluted in blocking buffer. At the end, they were counterstained for 5 min with Hoechst (2.5 μg/mL, Invitrogen, Cat# H3570) diluted in PBS at room temperature and washed 3 times for 5 min with PBS.

Images were acquired with either an IX81 inverted microscope (Olympus) with a 10x air objective (UPLFL PH 10x / 0.30, Olympus) or a 60x oil objective (PLAPO O 60x / 1.40, Olympus, Fig. 7A,B), or with an IX83 inverted microscope, equipped with a spinning disk unit (CSU-X1 10,000 rpm, Yokogawa) and a 40x silicone oil objective (UPLSAPO S 40x / 1.25, Olympus) and an Orca-Flash 4.0 camera (Hamamatsu) with the Olympus CellSens Dimension 2.2 software. Same acquisition settings were used to take pictures of the 6xHisTag staining (recombinant Shh) in all conditions. Three independent experiments were performed with similar outcome.

### Statistical analyses

Statistical analyses were performed with GraphPad Prism 7.02 software. All data were tested for normality (normal distribution) using the D’Agostino and Pearson omnibus K2 normality test and visual assessment of the normal quantile-quantile plot before choosing an appropriate (parametric or non-parametric) statistical test.

## Supporting information

spread sheet

## Author Contributions

A.D and N.W. performed experiments; A.D., N.W. and E.S. designed research, analyzed data and wrote the paper; K.L.T. contributed mice, reagents and critical comments on the paper.

## Acknowledgements

We thank James Briscoe, Raman Das, Michael Lin, Andrew McMahon, Thomas Thier and for constructs and probes. For mouse maintenance, genotyping and technical assistance, we are grateful to Beat Kunz, Marc Willaredt and Tiziana Flego. This work was supported by a grant from the Swiss National Science Foundation to E.S.. K.L.T. was supported by the Deutsche Forschungsgemeinschaft (DFG, SFB 488, Teilprojekt B9).

---

**Movie S1. Smo blockade with Cyclopamine induced aberrant dI1 axon guidance at the contralateral floorplate border.**

Twenty four hours time-lapse recording of the ventral midline of *ex vivo* spinal cords culture showing Math1::tdTomato-F-positive dI1 axons (black) at the contralateral floorplate border in an ethanol-treated control or Cyclopamine-treated sample. dI1 axons turned rostrally in an organized manner in the control, but many of them showed aberrant trajectories in the longitudinal axis in the presence of Cyclopamine. Maximum projections of z-stacks taken every 15 minutes are represented. EtOH, ethanol; Cyclop., Cyclopamine. Rostral is up.

**Movie S2. Smo blockade with Cyclopamine induced aberrant dI1 axon guidance at the contralateral floorplate border.**

Twenty four hours time-lapse recording of the ventral midline of *ex vivo* spinal cords showing Math1::tdTomato-F-positive dI1 axons (black) at the contralateral floorplate border in a Cyclopamine-treated sample. Axons that were crossing or about to exit the floorplate (filled arrowheads) at the beginning of the recording/inhibition showed a normal trajectory with a rostral turn. However, most of the axons that showed an aberrant guidance phenotype at the floorplate exit site (open arrowheads) were not yet in the floorplate at the time when the inhibitors were added to the medium. Cyclop., Cyclopamine. Rostral is up.

**Movie S3. Blockade of Smo entry into the cilium with Itraconazole induced aberrant dI1 axon guidance at the contralateral floorplate border**

Twenty four hours time-lapse recording of the ventral midline of *ex vivo* spinal cords showing Math1::tdTomato-F-positive dI1 axons (black) at the contralateral floorplate border in an DSMO-treated control or Itraconazole-treated sample. dI1 axons turned rostrally in an organized manner in the control, but many of them showed an aberrant trajectory in the longitudinal axis in the presence of Itraconazole. Maximum projections of z-stacks taken every 15 minutes are represented. Itracon., Itraconazole. Rostral is up.

