## Supplementary Figures for "A cell-autonomous role for primary cilia in long-range commissural axon guidance"

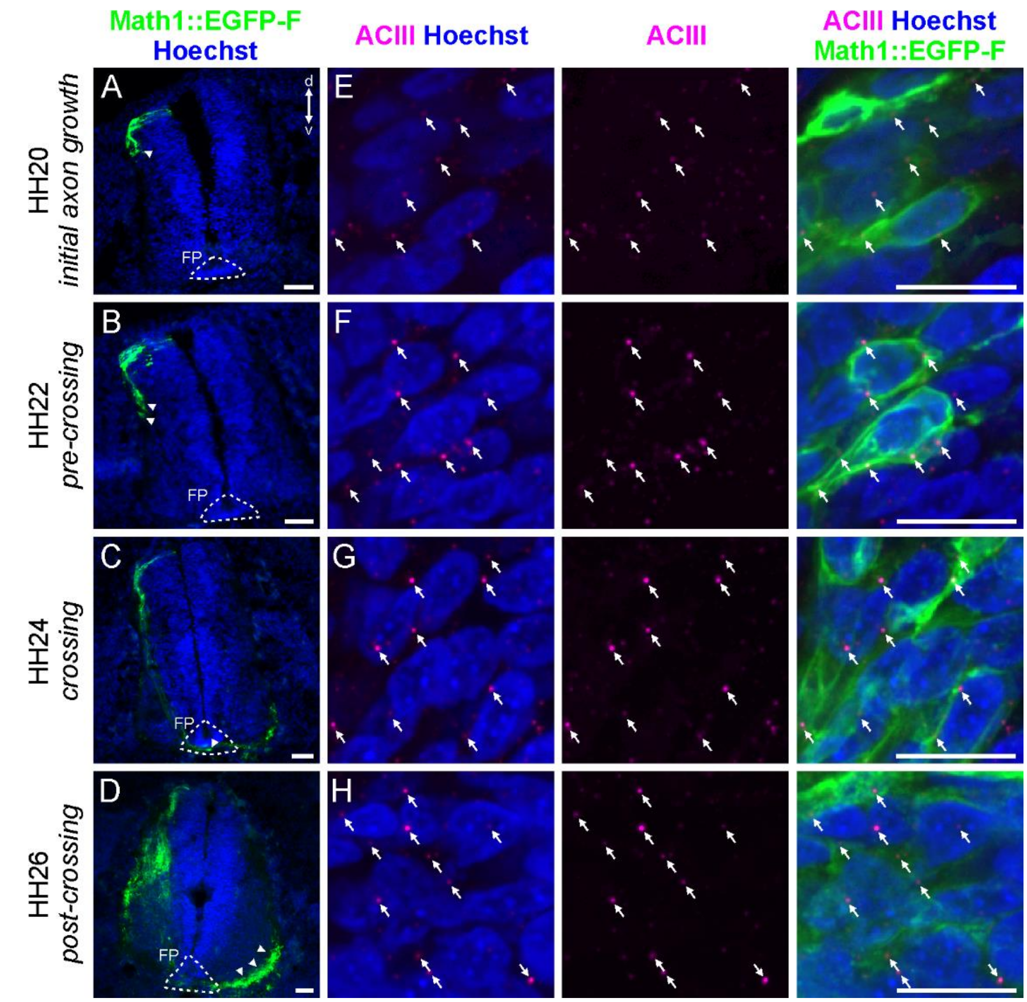


**Figure S1. ACIII staining confirms ciliation of dI1 commissural neurons during axonal navigation**. (A-D) Transverse sections of chicken embryos in which a Math1::EGFP-F plasmid (green) was electroporated unilaterally to label dI1 neurons at HH17-18. Embryos were sacrificed at different time points of their axonal navigation in the spinal cord. At HH20, dI1 axons were extending an axon (A). At HH22, they were growing ventrally but still pre-crossing (B). At HH24, they were crossing the floorplate (C) and at HH26, post-crossing axons were seen in the ventral funiculus (D). White arrowheads indicate where dI1 axonal growth cones localized at the different time points. Sections were counterstained with Hoechst to stain nuclei (blue). (E-H) High magnification pictures of the dI1 neuron area of spinal cords depicted in (A-D) showing the Math1-positive soma of dI1 neurons (expressing EGFP-F) co-stained with the primary cilium marker ACIII (magenta) and GFP (green). Thus, these neurons carried a primary cilium (white arrows) on their soma throughout stage HH20 to HH26. FP, floorplate; d, dorsal; v, ventral. Scale bars: 50 µm (A-D) and 10 µm (E-H).

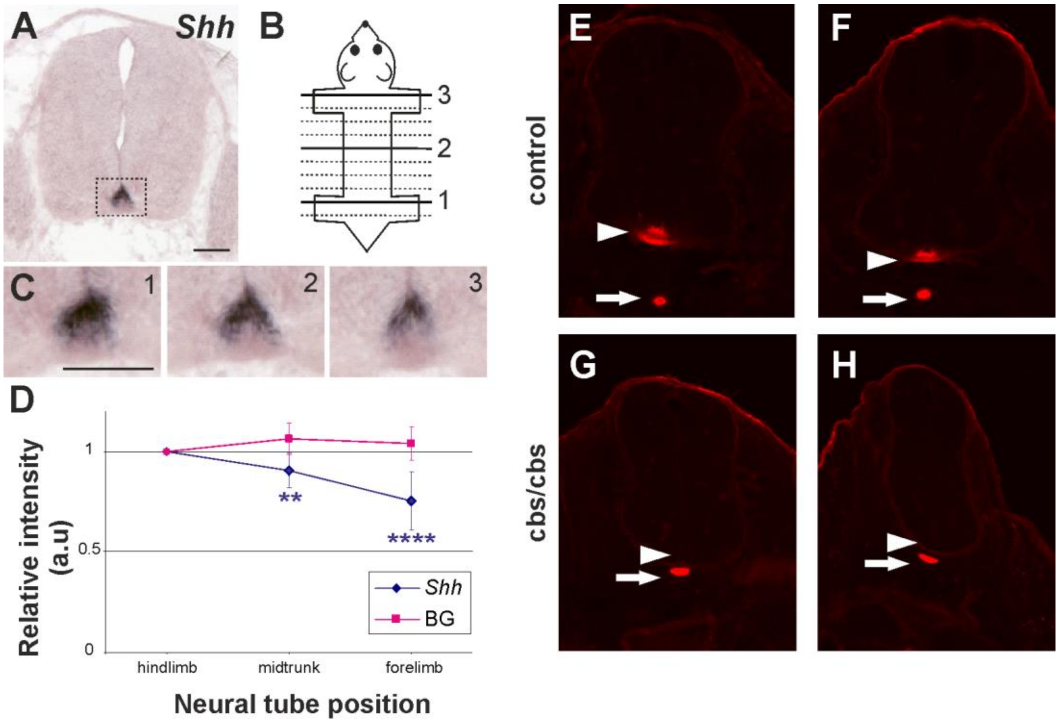


**Figure S2. Shh is expressed in an anterior to posterior gradient in the mouse spinal cord.**

(A) In situ hybridization for Shh shows specific expression in the floorplate (boxed area), in a transverse section of mouse E12.5 spinal cord. (B) Schematic illustration of the method used to collect sections spanning the longitudinal axis. Each slide contained 10 sections at 400 µm intervals. (C) Representative images of Shh in the floorplate at axial levels corresponding to hindlimb (1), midtrunk (2), and forelimb (3) of the same embryo. (D) Plot of relative Shh intensity versus relative position along the neural tube (mean ± SEM; n=12 embryos; single-sample T-test; $p<0.0001$ (****), $p<0.01$ (**)). Background (BG) staining levels, sampled from an area in the dorsal spinal cord, did not change along the A-P axis. a.u., arbitrary units. In wildtype embryos (+/+), Shh protein was also found in a decreasing gradient with higher levels in the caudal (arrowhead, E) and lower levels in the rostral floorplate (arrowhead, F). No or only very low levels of Shh were found in the floorplate of cbs mice (arrowhead, G,H). Shh was still expressed in the notochord of cbs mice (arrows). Scale bars: 100 µm. Source data and statistics are available in Source Data spreadsheet.

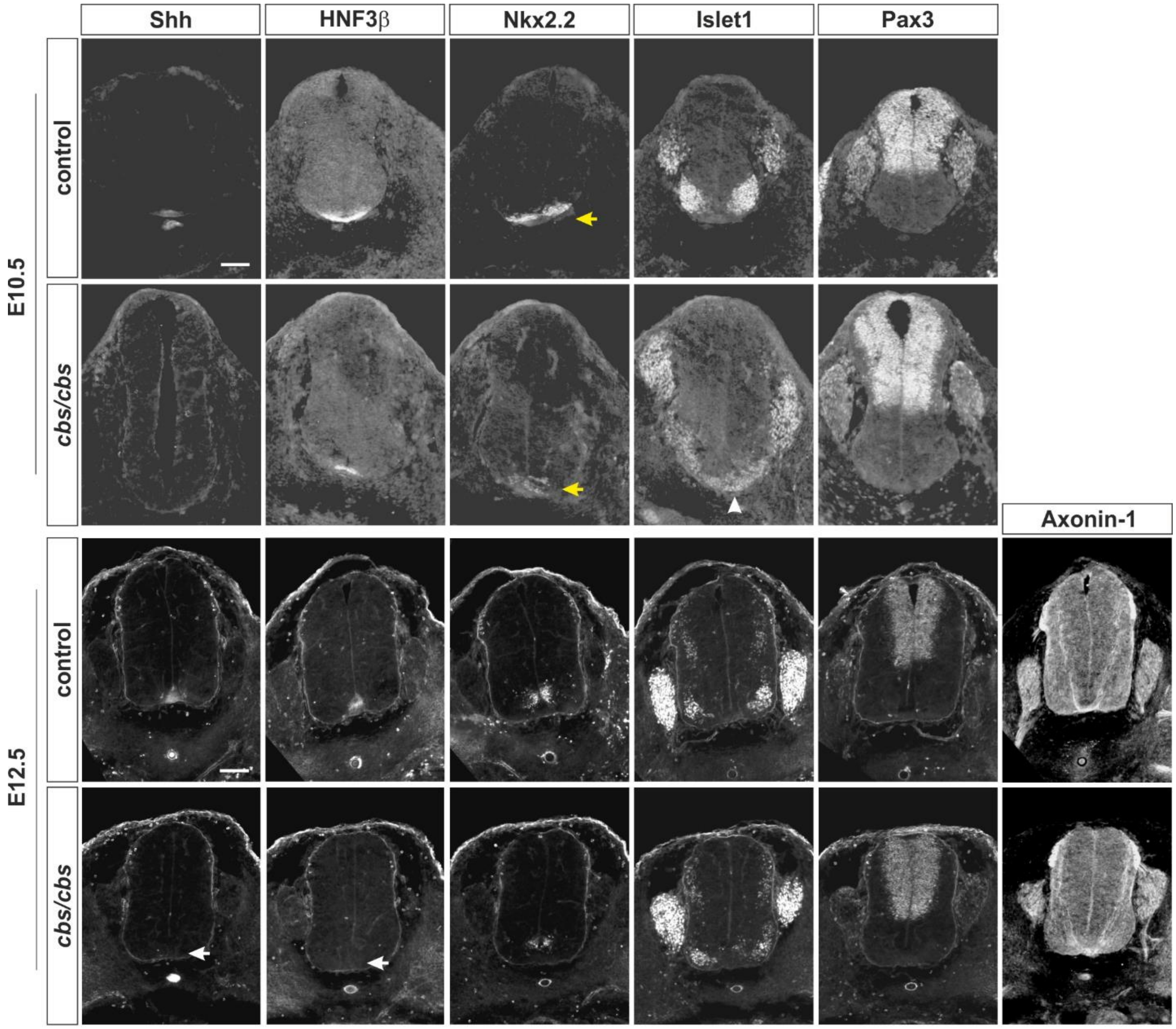


**Figure S3. Cbs mice display patterning defects in the ventral but not the dorsal spinal cord.** Immunostaining for an array of spinal cord markers (as indicated) at E10.5 and E12.5 revealed several defects in cbs embryos (arrows). Shh and the floorplate marker HNF3β were both missing from cbs spinal cords at E10.5 and E12.5. The ventral marker Nkx2.2 was present at both stages, but was reduced and disorganized, especially in E10.5 cbs mice. Islet1-positive cells erroneously invaded the ventral midline of E10.5 cbs mice, but by E12.5, Islet1 expression resembled that in control littermates. In contrast, the dorsal marker Pax3 was expressed normally in cbs mice. Dorsal commissural neurons, labeled by Axonin1/Contactin2, were also found in the normal position and projected correctly to the ventral spinal cord. However, the ventral commissure appeared defasciculated in cbs mice, as expected from our analysis of axon projections at the midline in open-book preparations. Scale bars: 50 µm.

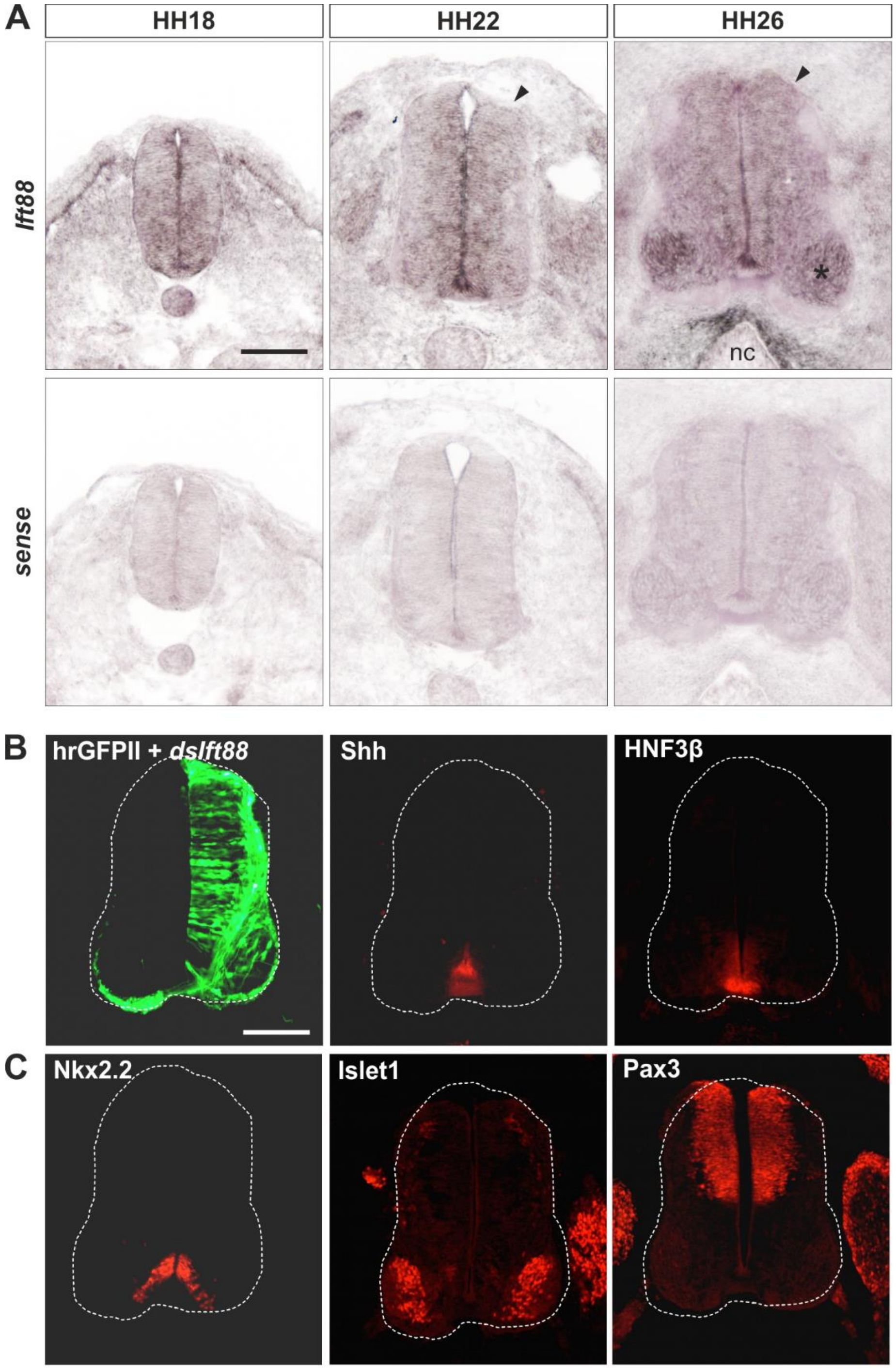


**Figure S4. Ift88 is expressed ubiquitously in the chick spinal cord, but its knockdown after HH18 does not affect early spinal cord patterning.**

(A) ISH for Ift88 at the indicated developmental ages (top) shows expression throughout the chick spinal cord, including the area occupied by the commissural neurons (arrowheads). At HH18 and

HH22, levels in precursors (ventricular zone) appeared to be higher than the expression in mature neurons. At HH26, Ift88 is expressed at slightly higher levels in the motoneurons (asterisk) but levels were no longer higher in the ventricular zone compared to the mantle zone. At this stage, Ift88 mRNA was also found surrounding the notochord (nc). No specific signal was seen with a sense control probe (bottom). (B,C) Immunostaining for a panel of spinal cord markers (as indicated) reveals normal patterning after electroporation of Ift88 dsRNA at HH18. The electroporated side was identified by expression of hrGFPII (green) from a co-electroporated plasmid. Downregulation of Ift88 at HH18, after spinal cord patterning is completed, did not affect Shh or HNF3β expression in the floorplate (B), nor affect patterning, as no difference in the expression of Nkx2.2, Islet1 or Pax3 were seen between the control and electroporated side (C). Scale bar 100 µm.

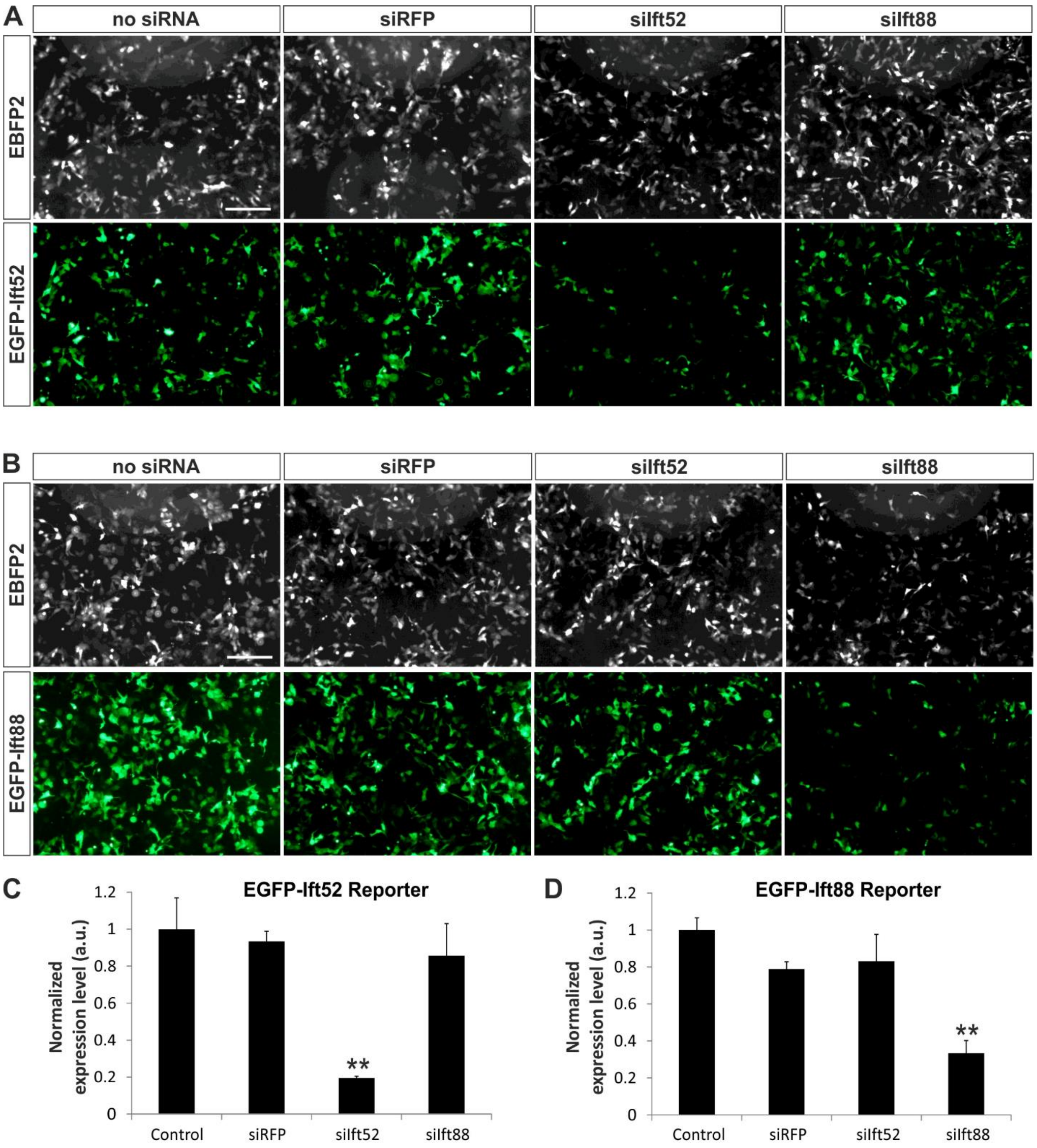


**Figure S5. Double-stranded RNAs against Ift88 and Ift52 knockdown target gene sequences efficiently and specifically.**

(A) COS7 cells were co-transfected with EBFP2 (top, transfection control), a reporter construct in which the Ift52 EST sequence was cloned downstream of EGFP (EGFP-Ift52; bottom) and siRNAs generated from long dsRNA, as indicated. EGFP expression was used to assess the efficiency and specificity of siRNAs against the Ift52 sequence. (B) COS7 cells were co-transfected as in panel (A), except that the reporter construct contained an Ift88 EST sequence cloned downstream of EGFP (EGFP-Ift88). (C-D) Quantifications. EGFP levels in each condition were normalized to EBFP2 transfection controls, and expression levels in the control (no siRNA) conditions were set to 1.0. (C) The EGFP-Ift52 reporter was reduced by 80.5±1.0% when co-transfected with siIft52 compared to the control condition. Co-transfection of siRFP or siIft88 did not significantly affect EGFP-Ift52 reporter levels. (D) The EGFP-Ift88 reporter was reduced by 66.7±6.9% when co-transfected with siIft88 compared to the control condition. Co-transfection of siRFP or siIft52 did not significantly affect EGFP-Ift88 reporter levels. n=10 measurements each; **$p<0.001$; One-way ANOVA with Tukey's multiple comparisons test. Scale bar 100 µm. Source data and statistics are available in Source Data spreadsheet.

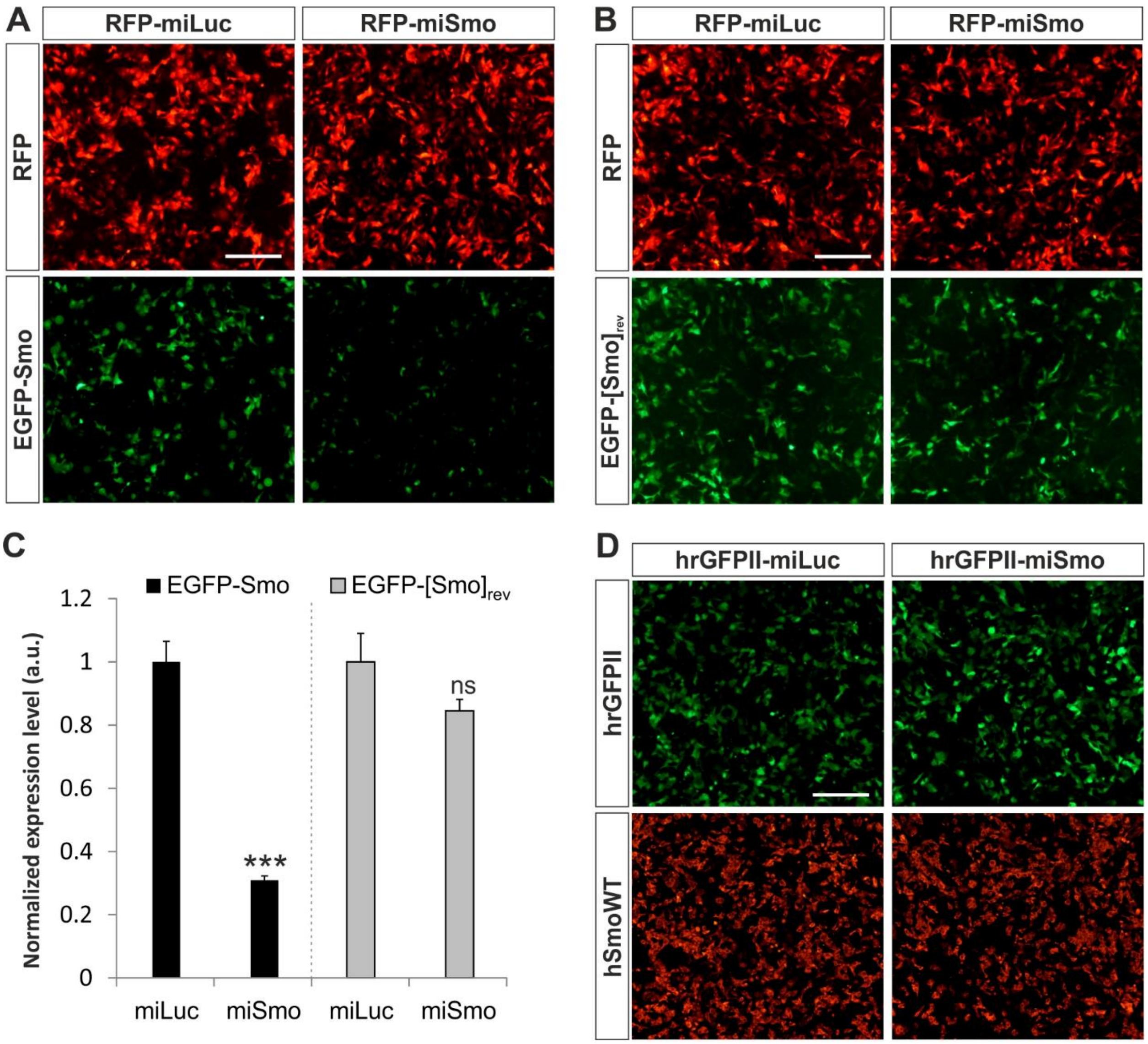


**Figure S6. Artificial miRNAs against Smoothened (miSmo) specifically and efficiently reduce target protein levels.**

(A) COS7 cells were co-transfected with pRFPRNAi vectors expressing artificial miRNAs against Luciferase (RFP-miLuc) or Smoothened (RFP-miSmo), together with a reporter construct in which a 2.2 kb fragment of chick Smo was cloned downstream of EGFP (EGFP-Smo). RFP expression (top) provided a transfection control, while EGFP expression (bottom) revealed the ability of the different miRNAs to knockdown the reporter gene. (B) COS7 cells were co-transfected as in panel

A, except the reporter construct contained a 2.2 kb fragment of chick Smo that was cloned in the reverse (antisense) orientation (EGFP-$[Smo]_{rev}$). (C) Quantifications. EGFP levels in each condition were normalized to RFP levels, and expression levels in the miLuc condition were set to 1.0. The EGFP-Smo reporter was reduced by 69.1±1.4% when co-transfected with miSmo compared to miLuc. In contrast, the EGFP-$[Smo]_{rev}$ reporter level was not significantly affected by miSmo transfection. n=15 measurements each; ***p<0.0001; Student's t-test. (D) COS7 cells were co-transfected with vectors expressing hrGFPII (green) and miRNAs against Luciferase (hrGFPII-miLuc) or chick Smoothened (hrGFPII-miSmo), together with a construct encoding human Smoothened (hSmoWT). Immunostaining for hSmoWT 24 hours after transfection (red) revealed that miSmo had no effect on hSmoWT expression levels compared to miLuc. Scale bars: 100 µm. Source data and statistics are available in Source Data spreadsheet.
